# QuickProt: A bioinformatics and visualization tool for DIA and PRM mass spectrometry-based proteomics datasets

**DOI:** 10.1101/2025.03.24.645047

**Authors:** Omar Arias-Gaguancela, Carmen Palii, Mehar Un Nissa, Marjorie Brand, Jeff Ranish

**Affiliations:** Institute for Systems Biology, Seattle, WA, USA; Wisconsin Blood Cancer Research Institute, Department of Cell and Regenerative Biology, Wisconsin Institutes for Medical Research, University of Wisconsin-Madison, Madison, WI, USA

**Keywords:** Data-independent acquisition, erythropoiesis, liquid chromatography-tandem mass spectrometry, mass spectrometry, parallel reaction monitoring, proteomics, QuickProt, stable isotope dilution

## Abstract

Mass spectrometry (MS)-based proteomics focuses on identifying and quantifying peptides and proteins in biological samples. Processing of MS-derived raw data, including deconvolution, alignment, and peptide-protein prediction, has been achieved through various software platforms. However, the downstream analysis, including quality control, visualizations, and interpretation of proteomics results remains challenging due to the lack of integrated tools to facilitate the analyses. To address this challenge, we developed QuickProt, a series of Python-based Google Colab notebooks for analyzing data-independent acquisition (DIA) and parallel reaction monitoring (PRM) proteomics datasets. These pipelines are designed so that users with no coding expertise can utilize the tool. Furthermore, as open-source code, QuickProt notebooks can be customized and incorporated into existing workflows. As proof of concept, we applied QuickProt to analyze in-house DIA and stable isotope dilution (SID)-PRM MS proteomics datasets from a time-course study of human erythropoiesis. The analysis resulted in annotated tables and publication-ready figures revealing a dynamic rearrangement of the proteome during erythroid differentiation, with the abundance of proteins linked to gene regulation, metabolic, and chromatin remodeling pathways increasing early in erythropoiesis. Altogether, these tools aim to automate and streamline DIA and PRM-MS proteomics data analysis, making it more efficient and less time-consuming.

## Introduction

Over the last few decades, mass spectrometry (MS)-based proteomics has emerged as one of the most dominant fields for protein profiling [1,2]. Advances in high-resolution MS instruments and computational tools have enabled researchers to characterize large-scale proteomes across various biological systems, with applications ranging from cell state transitions to structural biology and drug design [3,4]. Data-independent acquisition (DIA) is an untargeted MS technique used in proteomics analysis for the identification and relative quantification of peptides and proteins [5,6]. Unlike conventional approaches such as data-dependent acquisition (DDA), where precursor peptide ions are typically selected for fragmentation based on their intensities, DIA uses a series of predefined mass-to-charge (m/z) windows to select and fragment co-eluting precursor ions simultaneously [5,7,8]. This allows for the unbiased and often more reproducible identification of peptides and proteins, making it ideal for discovery proteomics experiments [9,10].

Parallel reaction monitoring (PRM) is a targeted MS strategy in which the mass spectrometer is programmed to analyze a predefined set of m/z values corresponding to peptides of interest [11,12]. By focusing on a predetermined set of target peptides, the technique can achieve low detection limits and high reproducibility levels. Compared to wide-window DIA, the resulting fragment ion chromatograms from PRM are less complex, simplifying their deconvolution and downstream processing [13,14]. Combined with stable isotope dilution (SID), PRM allows absolute quantification of peptides and proteins [15].

Although many software tools have been developed to identify and quantify peptide/proteins from raw LC-MS/MS data, such as DIA-NN [16], Spectronaut, OpenMS [17] and EncyclopeDIA [18], only a few offer the tools needed for quality control, data visualization, and statistical analysis (e.g., MSstats) [19] among the proteomes of experimental groups. While some tools, like Perseus [20], do provide some of these services, users are often limited to settings, pre-established by the developer, restricting the number and types of possible customizations. Here, we present QuickProt, a series of seven Python-based Google Colab notebooks dedicated to the data mining and visualization of DIA and PRM MS-proteomics datasets (Figure S1 and Table S1).

### QuickProt development and implementation

The QuickProt tool is designed to analyze DIA and PRM data through QuickProt-DIA and QuickProt-PRM modules, respectively (Figure 1, Figure S1). It enables proteomics data analysis to be more efficient and less time-consuming by automating several tasks. Users with no coding expertise can easily utilize these tools. The user only needs to provide a single input table generated from DIA-NN [16] or Skyline [21] software, two widely-used, freely available algorithms used for the initial analysis of DIA and PRM analysis, respectively. By simply clicking on the run button, users can execute all the codes in the notebooks. All outputs from these notebooks, including tables and figures, will be saved in dedicated folders. The resulting spreadsheet tables will be saved in a folder called ‘TABLES’ while the generated figures will be saved as TIFF images at a resolution of 300 dpi in the folder “PLOTS” (Table S1). Furthermore, as open-source code, these tools can be modified for other bioinformatics workflows depending on the needs of the investigator. All notebooks can be easily accessed at https://github.com/OmarArias-Gaguancela/QuickProt, where all of the modules have descriptions and links to their respective notebooks or pipelines.

**Figure 1:**
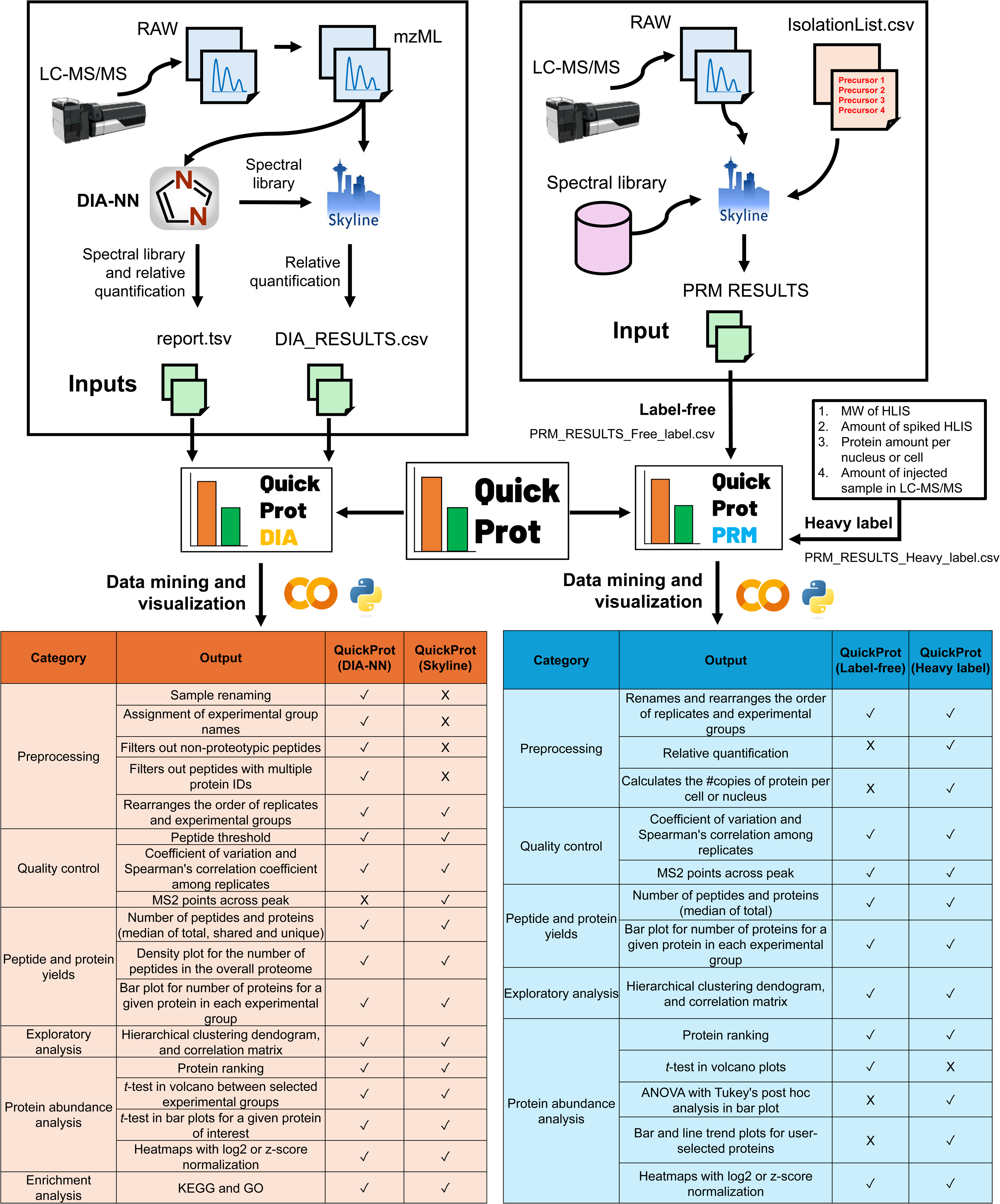
Overview of QuickProt Google Colab notebook tool for DIA and PRM analysis. In the left panel, the QuickProt-DIA workflow is represented. LC-MS/MS data is exported in RAW file format and subsequently converted to mzML format. Next, mzML files are imported into DIA-NN software for peptide and protein identification and relative quantification. Alternatively, mzML files can be imported into Skyline for quantification while using the DIA-NN spectral library for identification. Skyline or DIA-NN tables from either workflow can be analyzed in the Python-based QuickProt-DIA module composed of QuickProt-DIA (DIA-NN) or QuickProt (Skyline) notebooks. Both generate annotated tables and figures that fall into the following categories: preprocessing (e.g., sample filtering), quality control (e.g., coefficient of variation), peptide and protein yields, exploratory analysis (e.g., hierarchical clustering dendrogram), protein abundance (e.g., heatmap visualization), and enrichment analysis (e.g., KEGG). In the right panel, the QuickProt-PRM module workflow is represented. LC-MS/MS data in RAW file format is imported into Skyline alongside an isolation list (IsolationList.csv) and a selected spectral library. Following Skyline analysis, the output table ‘PRM_RESULTS_Free_label.csv’ or ‘PRM_RESULTS_Heaby_label.csv’ is imported into QuickProt-PRM for analysis. QuickProt (Label-free) notebook outputs are generated automatically, whereas, for heavy-label experiments, QuickProt (Heavy-label) notebook requires additional information (e.g., amount of spiked HLIS) for the generation of absolute quantities. The lower left and right panels show tables with the contents produced for each notebook. Abbreviations: LC-MS/MS: liquid chromatography-tandem mass spectrometry; mzML: Mass Spectrometry Markup Language; MW: molecular weight; HLIS: heavy labeled internal standard; KEGG: Kyoto Encyclopedia of Genes and Genomes; GO: gene ontology.

### QuickProt-DIA module overview

The QuickProt-DIA module includes two pipelines, depending on whether the input file is obtained from DIA-NN or Skyline. (Figure 1, Figures S1-S5). QuickProt-DIA (DIA-NN) pipeline uses the output table from DIA-NN, namely ‘report.tsv’, as an input file that contains peptides and proteins identified through their neural network-based spectral library and quantified by a MaxLFQ-like algorithm [16]. On the other hand, QuickProt-DIA (Skyline) pipeline uses a table from Skyline named ‘DIA_RESULTS.csv’ for processing. In QuickProt-DIA (Skyline), we used the DIA-NN spectral library for identification and Skyline to quantify peptides and proteins. Depending on the input table (DIA-NN or Skyline), the QuickProt-DIA module produces outputs that fall into the following categories: preprocessing, quality control, analysis of peptide and protein yields, exploratory analysis, protein abundance analysis, and gene enrichment analysis (Figure 1).

#### Preprocessing

During the preprocessing stage, the user can rename the samples and add experimental group names to the replicates of a given condition (Figure S2), a function not currently available in DIA-NN but provided in Skyline. In the case of QuickProt-DIA (DIA-NN), we filtered out non-proteotypic peptides and peptides that were assigned multiple Protein IDs in the DIA-NN prediction. Notably, both QuickProt-DIA pipelines allow customization of the layout of the samples and experimental groups. We incorporated an optional control point to filter out proteins that do not meet a specified unique peptide threshold (Figure S2). This feature is particularly beneficial for removing proteins that are identified by only a single peptide, which could result in false positives and/or unreliable quantification [22]. Implementing this type of quality control is recommended in discovery MS proteomics experiments [22].

#### Quality Control

For quality control purposes, we included a section to generate a coefficient of variation (CV) plot and correlation plot that displays Spearman’s correlation coefficient and data distribution among the replicates/samples of a given experimental group. For QuickProt-DIA (Skyline), we added a section for plotting the distribution of MS points across the chromatographic peaks for each experimental group. Since DIA quantification is generally based on MS2 points, we used MS2 points as the default choice. However, this can be adjusted to MS1, depending on the objective of the analysis (Figure S3). We highly recommend first running these sections of the notebook, as these outputs help to assess the reproducibility and technical variability in the experiment.

#### Peptide and Protein Yield

Next, QuickProt-DIA generates annotated tables and plots for the total number of peptides and proteins found in a given experiment (Figure S3). Additionally, it provides a comprehensive breakdown of the number of shared proteins between experimental groups and the number of unique proteins in each experimental group. These results are concurrently extracted into annotated CSV tables, which are deposited in a subfolder called SHARED_UNIQUE_PROTEINS within the TABLES folder, allowing easy access for the user (Table S1). These pipelines also allow the user to display a density plot for the number of peptides and calculate the median number of peptides per protein in the dataset. Additionally, users can enter the name of a specific protein or gene of interest to automatically visualize the number of peptides detected for that protein in the dataset.

#### Exploratory Analysis

In the exploratory analysis, the user can display hierarchical clustering dendrogram and correlation matrix plots to identify groupings and connectivity relationships between the experimental groups tested in the experiment (Figure S3).

#### Protein Abundance

For protein abundance visualization (Figure S4), we introduce a protein ranking tool where the user can array the proteins identified in each experimental group and rank them based on their estimated abundance levels. We provide a dedicated section for visualizing relative protein abundances through a volcano plot. For this analysis, a *t*-test is performed. Users can compare the fold change (FC) and *P*-values of the proteins between two experimental groups. To make it more interactive and user-friendly, we added drop-down menus allowing users to select *P*-values (0.05 to 0.001) and FC thresholds (0.5 to 10) that fit the experiment’s needs. These values are then converted to -log10 and log2 scales for plotting. Accordingly, a list of the total number of proteins analyzed is generated as an output table, along with lists of only upregulated or downregulated proteins in a subfolder called VOLCANO_PLOT_VALUES within the TABLES folder (Table S1). We went a step further by creating a section (Figure S4) that allows the user to visualize individual protein abundances across experimental samples by simply typing the name of the protein or gene of interest and then clicking a “Generate Plot” button to yield a bar plot with its respective statistical analysis.

The user selects the reference experimental group against which the statistical analysis will be performed. The bars of the plot will be automatically assigned an asterisk for statistically significant differences, or the letters ‘n.s.’ will be added on top of the bar when no statistical differences in protein abundances are found between the samples of interest. The user has the option to plot the abundances of all quantified proteins in each experimental group or sample via clustering heatmap visualization (Figure S4). We added a section where the user can input a list of proteins or genes of interest to be displayed in a heatmap. In this case, neither imputation nor clustering is conducted. Missing values are depicted with the letters ‘n.d,’ whereas the normalized values for the abundances of the selected proteins are shown on the log2 scale. Alternatively in another section, we used the ‘IterativeImputer’ tool from the Scikit-learn machine learning library [23], which uses an iterative approach to handle missing values, and then this data is used to display a clustering heatmap comparing the proteomes among experimental groups.

#### Enrichment Analysis

We incorporated the Kyoto Encyclopedia of Genes and Genomes (KEGG) and Gene Ontology (GO) tools in the enrichment analysis section of the QuickProt-DIA pipelines to identify biological pathways, processes, components, and functions that are enriched in a dataset. Hence, QuickProt-DIA facilitates KEGG and GO analyses on DIA datasets to provide a systems view of the biological processes that are affected in an experimental group, which can help prioritize proteins for potential follow-up experiments. In these pipelines, the user can choose whether to perform the analysis on the list of total proteins of a given experimental group, the total number of proteins compared in the volcano plot, or exclusively on upregulated or downregulated proteins derived from the same volcano plot analysis (Figure S5).

### QuickProt-PRM module overview

QuickProt-PRM module comprises two pipelines, namely, QuickProt-PRM (Label-free) and QuickProt-PRM (Heavy label), for processing of PRM data from Skyline through input tables titled ‘PRM_RESULTS_Free_label.csv’ or ‘PRM_RESULTS_Heavy_label.csv’, respectively (Figure 1, Figures S6-S10). Similar to the DIA module, QuickProt-PRM also produces outputs including categories such as preprocessing, quality control, peptide and protein yields, exploratory analysis, and protein abundance analysis. During the preprocessing step, customization of the sample and experimental group layout is provided for both notebooks (Figures S6-S7). QuickProt-PRM (Label-free) is dedicated to relative abundance quantification, whereas QuickProt-PRM (Heavy label) aims to calculate the number of molecules of a given protein in a sample using spiked-in isotopically heavy labeled internal standard (HLIS) proteins or peptides. These values can then be converted to copies per cell or organelle (e.g., nucleus) with the appropriate conversion factors. To accomplish such absolute calculations, the user must provide the molecular weight of the spiked HLIS, the known amount of protein per nucleus or cell used in the experiment, and the amount of sample injected into the LC-MS/MS (Figure S7). CV, correlation, and MS2 point plots can be generated during the quality control stage. The user can plot the number of quantified peptides for a given protein/gene, and view the total number of peptides and proteins quantified. Exploratory analysis through hierarchical clustering dendrogram and correlation matrix is also provided (Figure S8).

Like in QuickProt-DIA, QuickProt-PRM (Label-free) allows the user to visualize the estimated relative abundance of proteins via protein ranking, volcano, bar plots, and heatmaps (Figure S9). For QuickProt-PRM (Heavy label), absolute quantities of a protein (e.g., #copies/ nucleus or #copies/ cell) can be visualized through bar and/or line trend plots. We provide a section where users can select a given protein of interest and assess the statistical differences in abundance among experimental groups. In addition, one of the advantages of using PRM proteomics with HLIS is the ability to compare stoichiometric changes between different proteins. QuickProt-PRM leverages this by enabling an option to display and assess the statistical differences (Figure S10) between different proteins in a given experimental group. In both types of comparisons, ANOVA with Tukey’s post-hoc statistical tests are performed, and the results are displayed automatically in the bar plots. Indeed, we used the compact letter display to automatically depict significant differences among multiple groups. Finally, QuickProt-PRM allows users to display the abundance of all the targeted proteins for all experimental groups or by sample via clustering heatmaps. There is also an option to specifically display certain proteins of interest in the heatmap.

Lastly, we introduce two more modules, named QuickProt-PepSeq and QuickProt-ID Search. The first module is designed to evaluate the number of peptides identified in a specific amino acid sequence (e.g., domain) from a DIA dataset (Figure S11). This module is compatible with inputs from DIA-NN or Skyline reports processed using QuickProt-DIA, specifically ‘report_updated.csv’ or ‘DIA_RESULTS_UPDATED.csv’, respectively. In both notebooks, users simply need to input the sequence of interest and click the run button to produce an Excel spreadsheet (Table S1) with annotated tabs, displaying the number of peptide matches and their respective sequences by sample and experimental group. A bar plot of the number of peptide matches is also displayed (Figure S11). QuickProt-ID Search maps protein IDs in the UniProt database to a list of gene names provided by the user (Figure S12). This is especially helpful in cases where the investigator deals with thousands of genes.

### DIA dataset and analysis using QuickProt-DIA

#### DIA dataset

To demonstrate the performance of QuickProt-DIA, we analyzed an in-house generated DIA dataset. We employed a well-characterized *ex vivo* culture system method in which cord blood-derived human multipotent hematopoietic stem and progenitor cells (HSPCs) were induced to differentiate along the erythroid lineage [24,25] (Method S1). Samples were collected on days 0, 2, 4, 6, 8, 10, 11, 12, and 14, representing sequential stages of erythropoiesis, thus yielding cell lineages from HSPCs (day 0) to polychromatophilic and orthochromatic erythroblasts (day 14). Nuclear extracts were prepared from the cells (Method S1) and used to generate peptide samples for LC-MS/MS analysis on an Orbitrap Eclipse mass spectrometer operated in DIA mode (Method S2-S4; Table S2 and S4). During further processing, we used DIA-NN to generate a spectral library for peptide and protein identification due to its high sensitivity and processing speed. Peptide-to-protein quantification was achieved with Skyline. Chromatograms were extracted, and qualitative analysis of each product ion peak was performed, providing an additional safeguard to enhance data reliability. The quantitative information was exported from Skyline which was then processed using the QuickProt-DIA (Skyline) pipeline (Figure 2).

**Figure 2:**
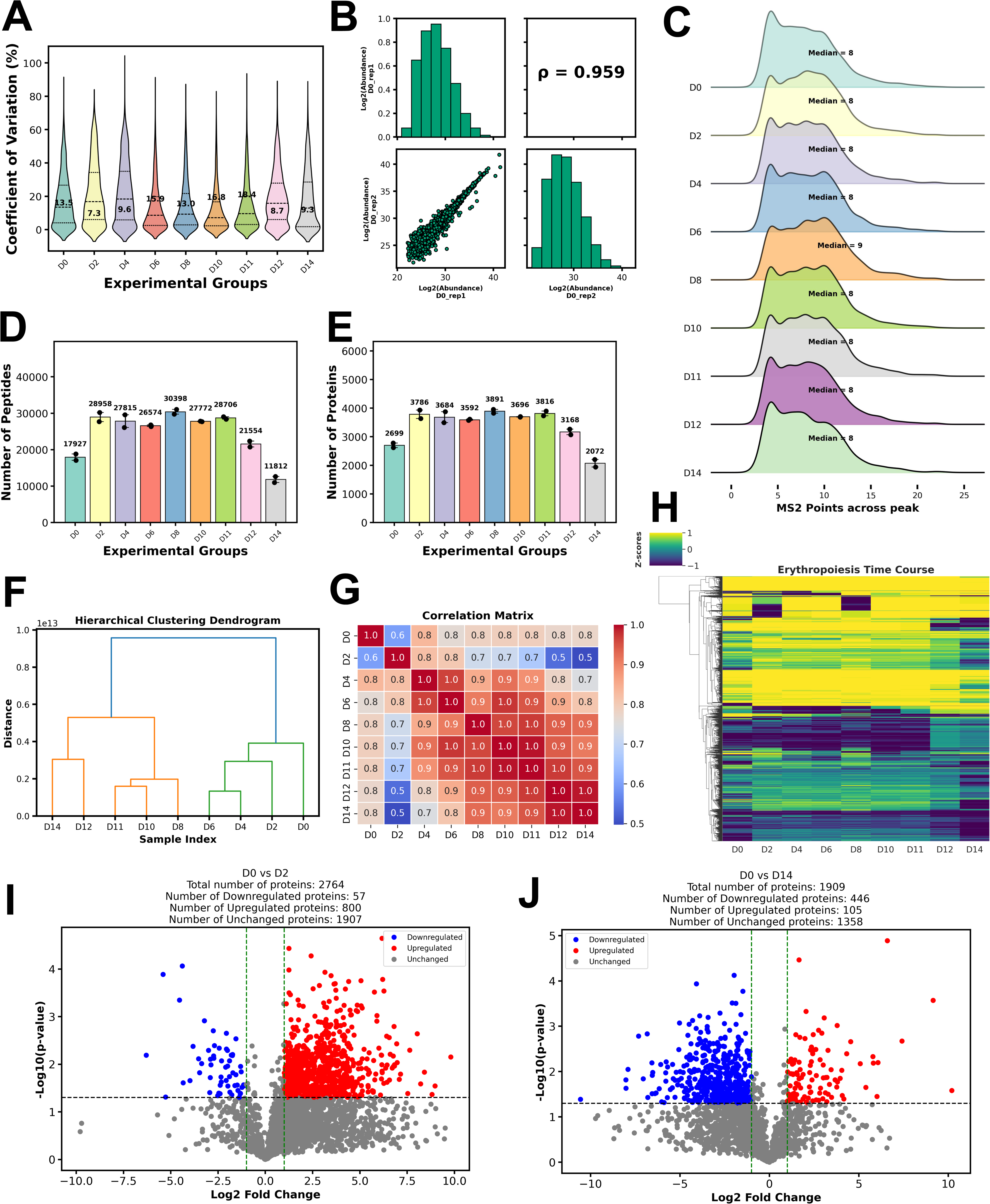
Comprehensive analysis of erythropoiesis time-course DIA proteomics data via QuickProt-DIA (Skyline) notebook. Samples on days 0, 2, 4, 6, 8, 10, 11, 12, and 14 (D0-14) during erythroid differentiation were collected for DIA discovery proteomics analysis. A) Coefficient of variation, with median values depicted inside each violin plot. B) Spearman’s correlation coefficient (ρ) and data distribution plots among replicates for day 0. C) Distribution of MS2 data points across the chromatographic peak, with median values depicted for the proteomes of each experimental group. D) Number of peptides identified in each experimental group, with median values depicted on top of each bar graph. E) Number of proteins identified in each experimental group, with median values depicted on top of each bar graph. Data represent the median ± SD of two biological replicates. F) Hierarchical clustering dendrogram. G) Correlation matrix among experimental groups. H) Clustering heatmap. I) Volcano plot for differential expression between D0 and D2. J), and D0 and D14. For I) and J), a *P*-value ≤ 0.05 and an FC (fold-change) threshold of > |2| were chosen for the analyses.

#### DIA analysis results

For this analysis, the minimum peptide threshold per protein was set at two for all samples in the dataset. Then, QuickProt-DIA was used to evaluate the quality and reproducibility of the DIA data. The CVs varied from 7.3% to 18.4% in the experimental groups tested (Figure 2A). Since these values were below the recommended 20% threshold [26–28], this suggests acceptable biological variability among the replicates. The correlation plot for Day 0 replicates shows a Spearman’s rank correlation coefficient of 0.959 (Figure 2B), and similar results were obtained for the other days of the time course (Figure S13). We also examined the distribution of MS2 points across chromatographic peaks; in this experiment, the median ranged from 8 to 9 MS2 points (Figure 2C). This range has been reported to be optimal for appropriately representing the peptide peak shape for accurate quantification [29]. Lastly, we assessed the median number of peptides per protein in each experimental group and found a median value of 4 to 5 for peptides per protein (Figure S14A). These metrics suggest that the LC-MS/MS method used was able to meet certain quality control requirements, ensuring robust quantitative accuracy for the experiment.

Regarding peptide and protein identification during the time course study, the total number of peptides varied from 11,812 to 30,398 (Figure 2D), yielding 2,072 to 3,891 proteins (Figure 2E), respectively. Notably, the core proteome representing the number of proteins shared between all experimental groups consisted of 1,813 identified proteins. The number of shared proteins between different samples varied from 96 to 1,958 (Figure S14B). Day 8 had the highest number of unique proteins (75), and Days 6 and 12 had the lowest number of unique proteins (3-4) (Figure S14C).

Inspection of the outputs from the exploratory analysis tools revealed a marked grouping at certain stages of erythropoiesis progression. Hierarchical clustering identified two major branches. The first branch comprised two sub-branches: one with day 0, and the other with days 2, 4, and 6. By contrast, the second branch contained two major sub-branches; one comprised of days 12 and 14, and the other of days 8, 10, and 11 (Figure 2F). In the correlation matrix, days 0 and 2 had the lowest levels of correlation compared to the other days of the time course, particularly when compared to days 12 and 14 (Figure 2G). This suggests timely and dynamic changes from early to late stages of erythropoiesis. This pattern is reminiscent of what was observed in the clustering analysis. When performing a clustering heatmap, the protein expression profiles for each day were readily visualized, and differentially expressed proteins could be easily detected. (Figure 2H). Given the important role of chromatin-modifying complexes in transcriptional regulation during erythropoiesis through modulation of chromatin accessibility [30], we used QuickProt-DIA to inspect the expression profiles of subunits of a few chromatin-modifying complexes [31], including the following: BAF (BRG1- or BRM-associated factor), ISWI (Imitation Switch), NuRD (Nucleosome Remodeling and Deacetylase), INO80 (Inositol requiring 80), SAGA (Spt-Ada-Gcn5 Acetyltransferase), ATAC (Ada Two-A Containing), SRCAP (Snf2-related CREBBP Activator Protein), and ATR-X (Alpha Thalassemia/Mental Retardation Syndrome X-linked) (Figure S15). QuickProt-DIA generated heatmaps that allowed easy visualization of the expression patterns and relative abundances for the members of these complexes over the time course. Interestingly, the abundances of several members of the BAF, e.g., ARID1A, and ISWI, e.g., SMARCA5 complexes, peaked at days 2 and 8, implying a requirement for these subunits at these specific time points during erythropoiesis (Figure S15).

Next, we used QuickProt-DIA to evaluate the protein abundance rankings for proteomes on each day of erythropoiesis (Figure S16). Notably, AHNAK was consistently the most abundant protein at all-time points.

Supported by the initial evidence from the exploratory analysis, we hypothesized that the number of upregulated proteins would increase significantly from the initial starting point on day 0 throughout certain time points during erythropoiesis progression. To validate this hypothesis, we used the volcano plot tool (Figure 2I-J; Figure S17) in the QuickProt-DIA (Skyline) pipeline and performed a statistical analysis of the proteomes of day 0 compared to each subsequent day. Similar to the day 0 vs. day 2 comparison (Figure 2I), when day 0 was compared to days 4-11, approximately 93% of the significantly changed proteins were upregulated, while the rest were downregulated. Notably, after day 11, the number of upregulated proteins declined, continuing until day 14 (Figure 2I and J; Figure S17). On day 14, out of the 551 significantly changed proteins, around 81% were downregulated, and 19% were upregulated (Figure 2J). Consistent with these findings, when inspecting individual proteins (e.g., ARID1A and SMARCC2), we observed a dramatic spike on day 2 and a decrease on days 12 and 14 compared to day 0 (Figure S18). These data reveal the dynamics of the proteome during erythropoiesis, and suggest that specific proteins may need to be upregulated in the early stages, whereas certain proteins may require downregulation in the later stages for proper differentiation

Lastly, we used the KEGG and GO tools in the QuickProt-DIA notebook to evaluate the biological role of proteins identified during different stages of erythropoiesis (Figures S19-S26). KEGG and GO data showed no major differences when inspecting the total number of proteins (Figure S19, S22-S24). However, focusing on significantly upregulated or downregulated proteins revealed some interesting findings. For instance, in the day 0 vs. day 2 comparison, upregulated proteins at day 2 were enriched in processes like spliceosome activity, primary metabolism (e.g., Fatty acid metabolism), and ATP-dependent chromatin remodeling (Figure S20A). The enrichment of proteins involved in chromatin remodeling at day 2 is consistent with the notion of priming at the early stages of development when the chromatin is relatively open [32], but transcription is still low. Conversely, in the day 0 vs. day 14 comparison, pathways associated with phagosome vesicle production were most enriched (Figure S20B) at day 14, likely linked to the extrusion of nuclei in the late stages of erythropoiesis [33–35]. By contrast, downregulated proteins in the day 0 vs. day 2 comparison exclusively affected pyruvate and glycolysis metabolism, while the day 0 vs. day 14 comparison showed downregulation in pathways related to spliceosome activity, DNA replication, nucleocytoplasmic transport, carbon metabolism, among others (Figure S21). This analysis indicates that certain pathways enriched on day 2 are downregulated at later stages of erythroid development, suggesting a timely and dynamic regulation during erythropoiesis.

We assessed molecular function, cellular components, and biological processes in the GO analysis (Figures S22-S26). For example, the upregulated proteins on day 2 showed enrichment for translation, ribosome biogenesis, nucleic acid, and protein binding processes (Figure S25). At day 14, cellular and protein localization, transport and binding, and ATP-related processes were prominent (Figure S25). Conversely, downregulated proteins on day 2 favored processes such as hemoglobin binding and cellular detoxification, while on day 14, protein and mRNA binding and processing, and chromosome reorganization were affected (Figure S26). Altogether, the QuickProt DIA (Skyline) pipeline efficiently provides an overview of the biological processes underlying erythroid differentiation, establishing an initial framework for evaluating complex proteomic datasets to extract meaningful biological insights.

### PRM dataset and analysis using QuickProt-PRM

#### PRM dataset

The mammalian SWI/SNF complex, also known as the BAF complex, is an ATP-dependent chromatin remodeling complex that plays important roles in gene regulation during cell differentiation [36–42]. This remodeler is organized into three distinct configurations: canonical-BAF (cBAF), polybromo-associated-BAF (PBAF), and non-canonical-BAF (ncBAF) [36,43]. Each complex comprises 12–15 subunits, with some subunits shared between different complexes, and paralogous subunits present for several of them. Mutations in protein components of BAF have been linked to numerous forms of cancer, neurodevelopmental disorders, and defective erythropoiesis [36,37,40,44,45]. An analysis of BAF subunit stoichiometry during erythropoiesis would provide insights into the dynamic regulation of the complex during erythropoiesis and can add to the existing knowledge on its role in chromatin remodeling and transcription [40]. Towards this aim, we determined the absolute abundances of 21 subunits (including paralogues) of the cBAF complex (all subunits except for ACTB) by SID-PRM mass spectrometry during the time course of erythropoiesis. This dataset was then analyzed using QuickProt-PRM (Heavy label) for easy visualization and interpretation. For this data generation, we created heavily labeled concatemer proteins (QconCATs) [46,47] containing up to 4 peptides for each cBAF subunit. Known amounts of these QconCATs were then spiked into the nuclear extract fractions followed by enzymatic digestion (Method S2-3; Table S3-S7). Peptides obtained after co-digestion were cleaned, and analyzed by PRM-MS to acquire targeted spectral information for the cBAF subunits (Method S3-4; Table S3-S4). Raw data (Table S4) was then processed in Skyline, where, after chromatogram extraction, each product ion peak was qualitatively monitored to ensure the reliability of the corresponding quantitative information. After analysis and refinement, the values were exported and analyzed using the QuickProt-PRM (Heavy label) pipeline (Figure 3).

**Figure 3:**
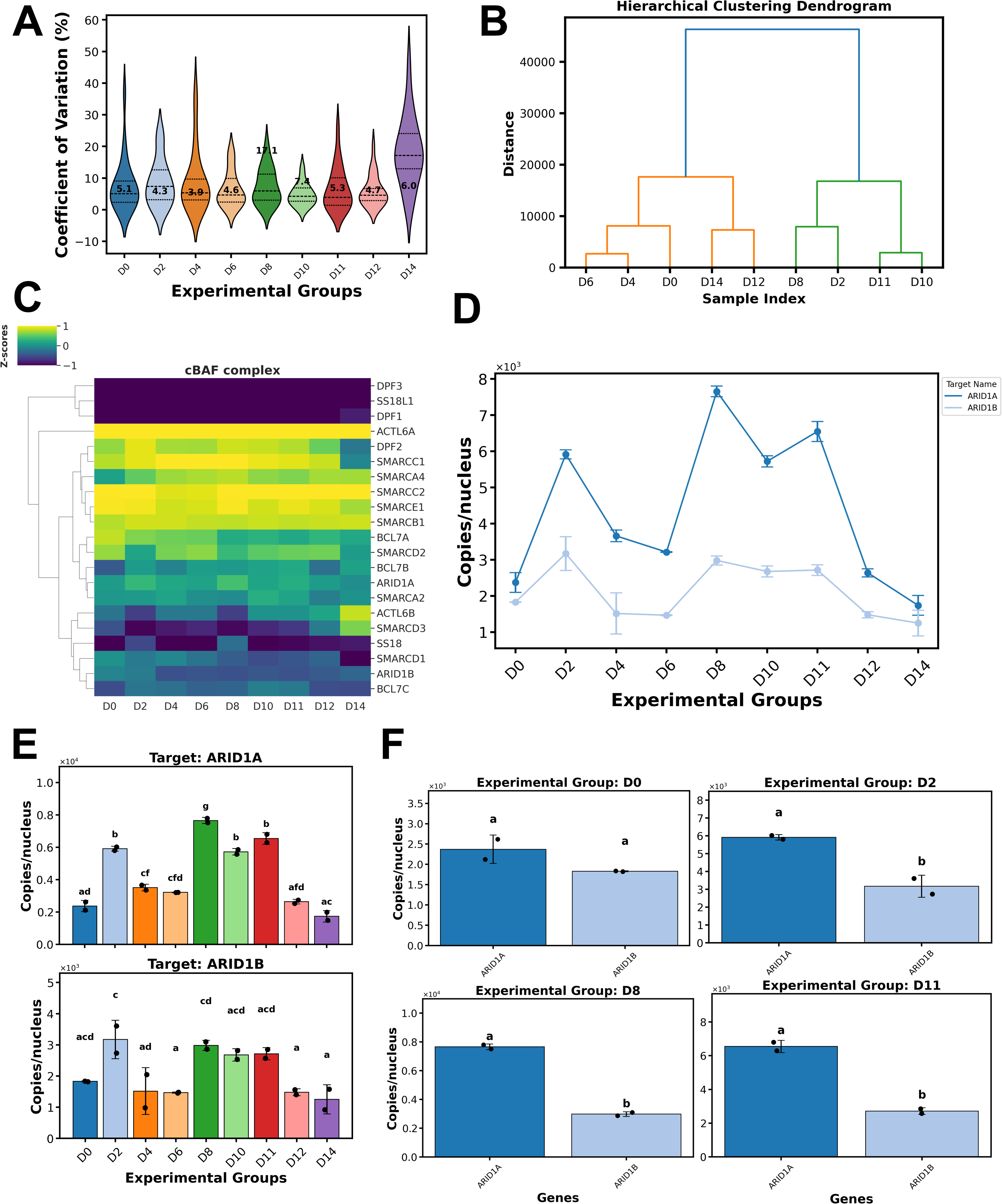
Targeted analysis of cBAF proteins during erythropoiesis via QuickProt-PRM (Heavy-label) notebook. Erythroid samples were spiked with heavily labeled concatamers (QconCATs) for absolute quantification (#copies/nucleus) of 21 cBAF proteins during erythroid differentiation via PRM proteomics. A) Coefficient of variation, with median values depicted inside each violin plot. B) Hierarchical clustering dendrogram. C) Clustering heatmap for proteins in the cBAF complex. D) Trend line plot for ARID1 paralogues, ARID1A and ARID1B. E) Bar plots for the absolute abundance of ARID1A and ARID1B, each plotted independently throughout the time course. F) Stoichiometric evaluation between ARID1A and ARID1B was conducted at four time points (D0, D2, D8, and D11). Data represent the median ± SD of two biological replicates. A compact letter display is automatically assigned on top of every bar. Different letters denote significant differences (*P* ≤ 0.05) by ANOVA with Tukey’s post-hoc test.

#### PRM analysis results

Quality control analysis revealed CVs ranging from 4.3-17.1% (Figure 3A). A Spearman’s rank correlation coefficient of approximately 0.9 among the samples (Figure S27) and a median of 28-32 MS2 points across chromatographic peaks (Figure S28A) was observed. These data indicate excellent reproducibility among the replicates from each experimental group. Exploratory analysis revealed two major groupings; the first comprised days 0, 4, 6, 12, and 14, while the second included days 2, 8, 10, and 11 (Figure 3B). Inspection of the correlation matrix showed high-level similarity among days, except for day 14, which had the lowest level of correlation compared to the others (Figure S28B).

Employing the peptide and protein yield option in QuickProt PRM (Heavy label), a total median peptide yield of 50-54 was detected for up to 21 cBAF proteins (Figure S29A-B). Peptide distribution showed a median of 2-3 peptides per protein detected (Figure S29C). Heatmap clustering and protein abundance ranking analysis revealed that among the 21 proteins quantified, ACTL6A was consistently the most abundant (42203 copies), whereas DPF1, DPF3, and SSL18L1 were the least abundant (137 copies) (Figure 3C; Figure S30; Table S8). Further, we inspected the relative stoichiometry of all cBAF paralogues during the erythroid time course (Figure 3D; Figure S31). For example, ARID1 paralogues, namely ARID1A and ARID1B, showed a similar number of copies per nucleus at day 0 (2,371 and 1,828 copies, respectively; *P*>0.05). Then, rapid and significant spikes in abundance were noticed on days 2, 8, and 11, followed by a decline until day 14 for both paralogues (Figure 3D-E). Noticeably, ARID1A yielded more copies per nucleus than ARID1B at most points of the time course (average difference of 2,763 copies on days 2-12; *P*≤0.05) (Figure 3D, F). Discordant expression levels between paralogues were also observed for ACTL6A over ACTL6B, DPF2 over DPF1 or DPF3, SMARCA4 over SMARCA2, and to a lesser extent in BCL7A over BCL7B and BCL7C, and SS18 over SS18L1 (Figure S31). Interestingly, SMARCD2 had a major spike exclusively on day 11, then dramatically decreased until day 14. Although SMARCD2 was mostly elevated relative to the other paralogues (SMARCD1 and SMARCD3), it is important to note a shift in relative abundance on day 14, where SMARCD3 was higher than the other two paralogues (Figure S31). The observed changes in the relative stoichiometry for these paralogues during the time course could indicate subunit switching [48,49] in cBAF or PBAF during erythroid development.

Unlike the other paralogues where the higher expression of one paralogue over another is evident, SMARCC1 and SMARCC2 yielded similar copy numbers throughout several data points, with some exceptions (e.g., day 14) (Figure S31). Lastly, to complete the analysis of cBAF members, we also analyzed the abundance trend of SMARCE1 (no paralogue) and found abundance spikes on days 2, 8, and 11 (Figure S31).

The data mining and visualization tools from the QuickProt-PRM notebook allowed us to efficiently assess the absolute quantities of members of the cBAF complex, highlighting their expression characteristics during erythropoiesis. This framework will be used in follow-up experiments to target specific cBAF subunits for knockdown or overexpression experiments to test the functional importance of the observed abundance measurements during erythroid differentiation.

## Conclusions and Outlook

QuickProt provides tools carefully designed to analyze and visualize data from DIA and PRM MS-proteomics experiments. In this article, we demonstrated the utility of the developed tools using in-house generated DIA and PRM datasets, offering insights into the biological processes underlying erythropoiesis. QuickProt offers comprehensive solutions beyond DIA data analysis (QuickProt-DIA). We provide notebooks for the analysis of PRM data (QuickProt-PRM), supporting both label-free and heavy-label experiments for relative and absolute quantification, respectively. Moreover, we have incorporated two more tools; QuickProt-PepSeq to report the number of unique peptide matches to a user-defined protein sequence in DIA datasets, and QuickProt-ID Search to retrieve UniProt protein IDs from gene names. We leveraged the accessibility of Google Colab notebooks to build our Python-based code within them. This makes it more user-friendly and versatile compared to other tools in the field, which typically require some coding skills. Furthermore, as an open-source platform, users can modify and customize the tools to suit the needs of specific projects.

## Acknowledgments

This work was supported by the National Institutes of Health (NIH), Grant number: RO1DK098449.

## Conflict of Interest Statement

The authors declare no conflicts of interest.

## Data availability Statement

The LC-MS/MS raw data, spectral libraries, and the input files processed in QuickProt have been deposited in ProteomeXchange [PXD060333] and can be accessed also via Panorama Public [https://panoramaweb.org/QuickProt_datasets.url]. The QuickProt notebooks were deposited in GitHub (https://github.com/OmarAriasGaguancela/QuickProt).

## Supplementary Methods

### Method S1: Hematopoietic stem and progenitor cell isolation, culture, and total protein extraction

Hematopoietic stem and progenitor cell isolation was performed as previously reported [15]. Briefly, CD34^+^ cells from umbilical cord blood donors were enriched by negative selection using the RosetteSep kit (STEMCELL Technologies; Catalog number: 15631) and the Ficoll density gradient, followed by positive selection with the Positive Selection Kit (STEMCELL Technologies, Catalog number: 18096). The purified cells were analyzed for CD34 expression via FACS (Fluorescence-activated cell sorting). CD34^+^ cells were differentiated into the erythroid lineage using a 4-step induction protocol. First, between days 0-11, CD34+ cells grew in a serum-free IMDM (Iscove′s Modified Dulbecco′s Medium) medium supplemented with various combinations of human recombinant cytokines and other additives. Second, between days 11-15, cells were co-cultured with stromal MS-5 cells in an enriched IMDM medium supplemented with EPO (erythropoietin). Third, between days 15-18, co-culture was transferred to the enriched IMDM medium without cytokines. Fourth, between days 18-26, co-culture was incubated in an enriched IMDM medium with 10% fetal bovine serum. Cell viability and counting were conducted by FACS analysis, with samples collected on days 0, 4, 6, 8, 10, 11, 12, and 14. Two biological replicates were collected for subsequent experiments.

For total nuclear protein extraction, 10 × 10^6^ cells were centrifuged at 1500 rpm for 5 minutes at 4°C. The cells were then washed with 1 mL of cold PBS (phosphate-buffered saline) pre-chilled to 4°C. The cells were suspended in 80 µL of swelling buffer (10 mM HEPES pH 8, 1.5 mM MgCl_2_, 10 mM KCl, 0.1% NP40, 1 tablet of protease inhibitor C per 10 mL of buffer), vortexed, and kept on ice for 15 minutes with vortexing every 5 minutes. Pellet nuclei were obtained by centrifugation at 1500 rpm for 5 minutes at 4°C. The nuclei pellets were then washed twice with 1 ml of ice-cold PBS and re-suspended in 80 µL of extraction buffer 1 (50 mM HEPES pH 8, 1.0 mM MgCl_2_, 150 mM NaCl, 1% NP40, 1 tablet of protease inhibitor C per 10 mL), vortexed 5 times for 2 seconds each, and centrifuged for 10 minutes at 13,000 rpm at 4°C. The supernatant, or nuclear extract 1 (NE1), was kept on ice or stored at −80°C. Next, 80 µL of extraction buffer 2 (50 mM HEPES pH 8, 1.0 mM MgCl_2_, 150 mM NaCl, 0.5% (w/v) sodium deoxycholate, 1% NP40, 0.1% (w/v) SDS, 50 ng/µL benzonase, 1 tablet of protease inhibitor C per 10 mL of buffer) was added to the pellet nuclei. The mixture was vortexed for 5 minutes, shaken on a thermomixer at 14,000 rpm and 37°C for 20 minutes, vortexed again for 5 minutes, and centrifuged for 15 minutes at 13,000 rpm at 4°C. The supernatant, or nuclear extract 2 (NE2), was kept on ice or stored at −80°C. Aliquots of 3 µL were taken from NE1 and NE2 for BCA protein assay measurement (Thermo Scientific; J63283.QA), following the manufacturer’s instructions. NE1 and NE2 were mixed and used for subsequent experiments.

### Method S2: Production of heavily labeled QconCATs

QconCATs (QconCAT09-12, Table S5) were cloned into the pEU-EO1-MCS expression vector (Cell Free Sciences). The coding sequences for each QconCAT are flanked by an N-terminal linker (MGASGK) and a Strep II tag (WSHPQFEK), followed by a C-terminal linker connected to a His-tag (YGGGSHHHHHHHH). Protein expression was conducted using a wheat germ cell-free protein expression system (WEPRO8240H kit; Cell Free Sciences) following the instructions of the manufacturer. During translation, isotopically heavy arginine (¹³C₆,¹⁵N_4_) and lysine (¹³C₆,¹⁵N₂) are incorporated in the QconCATs. His-tagged QconCAT proteins were purified via Ni-NTA (Nitrilotriacetic acid) chromatography. Briefly, 420 µL of binding buffer (30 mM Tris pH 8.0, 500 mM NaCl, 1 mM TCEP, 8 M urea, 10 mM imidazole) were mixed with a 210 µL aliquot from the QconCAT protocol. The mixture was incubated at room temperature for 30 minutes with end-over-end rotation. Separately, the Ni-NTA beads (Thermo Fisher; Catalog number: 88222) were mixed by tilting the bottle on a longitudinal axis until homogeneous. Then, 100 µL of Ni-NTA were taken into a 1.5 mL tube and washed with ∼1 mL of binding buffer. This was repeated twice, followed by centrifugation at RT, 700 g for 1 minute. The QconCAT sample was added to the washed Ni-NTA beads and incubated at RT for 60 minutes with end-over-end rotation. To equilibrate the spin column (Thermo Fisher; Catalog number: 89879), 400 µL of binding buffer was flowed through the column by centrifugation at 700xg for 1 minute at RT, and the flow-through was discarded. The sample and Ni-NTA beads were poured into the spin column inserted in a collection tube and centrifuged at 700xg, RT for 1 minute to collect the flow-through. The column was then inserted into a new collection tube and washed with 400 µL of wash buffer (30 Tris pH 8.0, 500mM NaCl, 1mM TCEP, 8 M urea, 25 Imidazole) three times. The column was moved to a new collection tube and eluted with 20 µL of elution buffer (30 mM Tris pH 8.0, 500 mM NaCl, 1 mM TCEP, 8 M urea, 250 Imidazole) 4-5 times. The eluates (His-tagged purified QconCATs) were stored at −80 °C. To assess the yield of His-tagged purified QconCATs, we used a known amount of commercially available Strep II tag peptide (GenScript) and quantified QconCAT amount via SID-PRM MS-proteomics.

### Method S3: Sample processing for MS-proteomics

The total protein of nuclear extracts was normalized to 10 µg for all erythroid samples. 2.5 ng of each heavy-labeled QconCAT (QconCAT09-12, Table S5) was spiked into the nuclear extracts. LCMS-grade water (Thermo Scientific; Catalog number: 047146.K2) was added to reach a total starting volume of 30 µL. Next, 30 µL of 2x lysis buffer (10% SDS [Sodium dodecyl sulfate]; 100 mM ABC [Ammonium Bicarbonate]) was added to the sample in safe-lock protein low-bind 1.5 mL tubes (Fisher Scientific; Catalog number: 13698794). For reduction, 2.6 µL of 120 mM TCEP (Thermo Scientific; Catalog number: 77720) was added and incubated at 37 °C for 15 minutes without shaking. For alkylation, 2.6 µL of 500 mM CAA (Thermo Scientific; Catalog number: 148410050) was added, and the sample was incubated in the dark for 20 minutes at room temperature (RT) without shaking. The denaturation process was completed with 6.6 µL of 55% phosphoric acid, followed by vortexing for 5 minutes. Then, 430 µL of an in-house prepared S-buffer (90% Methanol; 100 mM Tris, pH 8.0) was added and vortexed for 5 minutes. The samples were then added to columns from the S-Trap micro MS sample prep kit (Protifi; Catalog number: C02-micro-80) and centrifuged at 4,000 × g for 30 seconds at RT. This process was repeated in ∼100 µL increments until the entire sample volume passed through the column, with flow-through being discarded. The columns were washed four times with 150 µL of S-trap buffer via centrifugation (4,000 × g for 30 seconds at RT) and then dried by centrifugation (4,000 × g for 30 seconds at RT). The S-trap micro spin columns were placed into 2.0 mL low-binding tubes. For digestion, 25 µL of 1 µg Trypsin-LysC mix (Thermo Scientific; Catalog number: A4100) in 50 mM ABC was added to the columns and incubated at 37 °C overnight. Samples were eluted with 20 µL of 50 mM ABC, 40 µL of 0.2% formic acid (FA), and 50% acetonitrile (ACN). Each solvent was added twice and centrifuged at 4,000 × g for 30 seconds at RT. The combined eluate was dried in a speed vacuum, and the digested peptides were re-suspended in 60 µL of 0.1% FA, representing 10 µg of peptides in a final volume of 60 µL.

### Method S4: DIA and PRM sample processing in Eclipse Orbitrap LC-MS/MS

A volume of 3 µL (0.5µg of light peptides and 0.125 ng of heavy labeled peptides) was injected into Eclipse Orbitrap LC-MS/MS instrument equipped with precolumn Acclaim PepMap 100 C18 HPLC Columns (Thermo Scientific; Catalog number: 164946) and analytical column EASY-Spray HPLC Column (Thermo Scientific; Catalog number: ES900). For DIA, the LC followed a gradient of 0-85 min (4-25% ACN), 5 min (25-30% ACN), 5 min (30-80% ACN), and 10 min (80% ACN). For PRM, the LC followed a gradient of 0-5 min (2-5% ACN), 40 min (5-32% ACN), 5 min (32-60% ACN), 5 min (60-85% ACN), and additional 10 min (85% ACN). For both DIA and PRM, a flow rate of 300 nL/min was used. MS parameters for DIA and PRM are described in detail in Tables S2 and S3, respectively.

For DIA data processing in DIA-NN, we first created a reference library by uploading FASTA sequences from the UniProt human proteome. These sequences were then subjected to *in silico* digestion in DIA-NN with the following settings: 0.01 FDR, precursor range 395-1005 m/z, fragment ion range 200-2000 m/z, precursor charge range of 1-4, peptide length of 7-30, protease Trypsin/P (max cleavages of 1), and enabled N-terminal methionine excision and cysteine carbamidomethylation modifications. Next, we uploaded the LC-MS/MS files into MSConvertGUI [50] software to convert the files from RAW to mzML format. Experimental samples in mzML format were loaded into DIA-NN, along with the FASTA sequences and the predicted reference library. The final outputs generated were the report-lib.parquet.skyline.speclib, report-lib.parquet, and report.tsv files. Both report-lib.parquet.skyline.speclib and report-lib.parquet are sample-specific libraries generated by DIA-NN, whereas report.tsv contains identification and peptide-protein quantification data for the analyzed samples.

For DIA data processing in Skyline, we first imported the report-lib.parquet.skyline.speclib library with default settings. Then, LC-MS/MS samples in mzML format were imported into Skyline. Product ions were configured to have the best six transitions. In peptide settings, we chose a peptide length of 7-30, protease Trypsin/P (max cleavages of 1), and enabled N-terminal methionine excision and cysteine carbamidomethylation modifications. In transition settings, we chose a fragment ion range of 200-2000 m/z, retention time filtering of scans within 5 minutes of MS/MS IDs, and a precursor charge range of 1-4. To assess associated proteins, we used the following settings: peptides must be unique to a single protein, find the minimal protein list that explains all peptides, and remove subset proteins. To import peak boundaries, we changed the column headings in report.tsv from File.Name, Modified.Sequence, RT.Start, and RT.Stop to FileName, PeptideModifiedSequence, MinStartTime, and MaxEndTime, respectively, as these are compatible with Skyline. Finally, DIA results were exported in a CSV report called “DIA_RESULTS.csv” for downstream analysis in QuickProt-DIA (Skyline) notebooks.

For PRM upstream processing, the transition list “IsolationList.csv” file was imported into the Skyline environment, and then LC-MS/MS samples in RAW format were imported. Next, we imported an in-house BAF library called “BAF_Prosit1.blib” into Skyline. This library was created with the inbuilt Prosit tool in Skyline choosing 27 as the collision energy value. Heavy isotope labels (^13^C and ^15^N) to the C_6_ and N_2_, and C_6_ and N_4_ of lysine and arginine residues, respectively, were added in the peptide settings interface. PRM results were exported in a CSV report called “PRM_RESULTS_Heavy_label.csv” for downstream analysis in QuickProt-PRM (Heavy label) notebooks.

## Supplementary Figures

**Figure S1: Scheme of the QuickProt tool for proteomic data mining and visualization. QuickProt comprises five modules: QuickProt-DIA, QuickProt-PRM, QuickProt-PepSeq, and QuickProt-ID Search.** QuickProt-DIA consists of two notebooks or pipelines: QuickProt-DIA (DIA-NN) and QuickProt-DIA (Skyline). Additionally, QuickProt-PRM includes two notebooks: QuickProt-PRM (Heavy label) and QuickProt-PRM (Label-free). QuickProt-PepSeq features two pipelines: QuickProt-PepSeq (DIA-NN) and QuickProt-PepSeq (Skyline). Lastly, QuickProt-ID Search has a notebook of the same name.

**Figure S2: Overview of the QuickProt-DIA (DIA-NN) or (Skyline) notebook data preprocessing interface.**

**Figure S3: Overview of the quality control, peptide and protein yields, and exploratory analysis interface in the QuickProt-DIA (DIA-NN) or (Skyline) notebooks.**

**Figure S4: Overview of the protein abundance analysis interface in the QuickProt-DIA (DIA-NN) or (Skyline) notebooks.**

**Figure S5: Overview of the enrichment analysis interface in the QuickProt-DIA (DIA-NN) or (Skyline) notebooks.**

**Figure S6: Overview of the data preprocessing interface in the QuickProt-PRM (Label-free) notebook.**

**Figure S7: Overview of the data preprocessing interface in the QuickProt-PRM (Heavy label) notebook.**

**Figure S8: Overview of the quality control, peptide and protein yields, and exploratory analysis interface in the QuickProt-PRM (Label-free) or (Heavy label) notebooks.**

**Figure S9: Overview of the protein abundance analysis interface in the QuickProt-PRM (Label-free) notebooks.**

**Figure S10: Overview of the protein abundance analysis interface in the QuickProt-PRM (Heavy label) notebooks.**

**Figure S11: Overview of QuickProt-PepSeq workflow.** Outputs from DIA-NN or Skyline, coupled with QuickProt-DIA, can be imported into the QuickProt-PepSeq notebook (part of the QuickProt notebook series). A given region of interest (e.g., a protein domain) is then entered into the notebook to be mapped against the DIA data of a specific experiment. As a result, an Excel file, ‘PeptideMatches.xlsx’ will be generated. This file contains, in different tabs, the number of peptide matches per sample and per experimental group, as well as their respective amino acid sequences. The notebook also includes an option for plotting a bar graph showing the number of peptide matches in each experimental group.

**Figure S12: Overview of QuickProt-ID Search workflow.** The user inputs a list of genes of interest into QuickProt-ID. Using the Unipressed (Uniprot REST) library, the notebook has been adapted to generate a CSV table with their respective protein IDs.

**Figure S13: Spearman’s correlation coefficient (ρ) and data distribution plots among replicates for days, 2, 4, 6, 8, 10, 11, 12, and 14 in DIA-data.**

**Figure S14: Distribution of the number of peptides per protein and bar plots for shared and unique proteins in DIA data.** A) The number of peptides per protein plot depicts the distribution of values as well as the median value for each experimental group. Bar plots depict the median values for B) shared and C) unique proteins. A summary table with the names and numbers of proteins is generated and stored in a subfolder called ‘SHARED_UNIQUE_PROTEINS’ within the ‘TABLES’ folder. Additionally, within the same subfolder, another folder called ‘Extracted’ is generated to extract and store tables listing the proteins that are shared or unique for a given comparison or experimental group.

**Figure S15: Heatmaps show the relative abundances of proteins in the BAF, ISWI, NuRD, INO80, SAGA, ATAC, SRCAP, and ARTX chromatin remodeling complexes during the time course in the DIA data.** Log2 normalization was applied to the abundance values for proteins from days 0 to 14. The abbreviation ‘n.d.’ stands for ‘not detected’, indicating values that were not found in the datasets.

**Figure S16: Protein ranking for erythroid samples collected on days 0, 2, 4, 6, 8, 10, 11, 12, and 14 in DIA data.** The Y-axis represents the log2 abundance of proteins in the proteomes of each experimental group, whereas the X-axis depicts the abundance ranking for proteins in a given proteome. The names of the maximum, median, and minimum-ranking proteins are shown in the plot.

**Figure S17: Volcano plots for differential expression of D0 vs. D4, D6, D8, D10, D11, or D12 from the DIA data.** The volcano plots depict a *P*-value ≤ 0.05 and a fold-change (FC) threshold of 1.0. In the upper part of the plot, beside the title, a detailed description of the total number of upregulated, downregulated, and non-significantly changed proteins is shown.

**Figure S18: Bar plots for protein abundance and peptide numbers for selected proteins in the DIA data of the erythroid samples.** A) The abundance of ARID1A (upper panel) and SMARCC2 (lower panel) proteins is displayed in bar plots. Statistical analysis by *t*-test was performed on the datasets, using Day 0 as the reference group for comparison. Asterisks represent statistically significant differences (*P* ≤ 0.05), whereas ‘n.s.’ indicates non-significant differences. These statistical representations were added automatically by the code in the QuickProt-DIA notebook. B) Number of peptides supporting the abundance estimate for ARID1A and SMARCC2 in each experimental group. Data represent the median ± SD of two biological replicates.

**Figure S19: KEGG enrichment analysis of proteomes from individual experimental groups (A-C), and for D0 vs. D2 or D14 comparisons (D-E) in DIA data.**

**Figure S20: KEGG enrichment analysis of upregulated proteins in proteomes for A) D0 vs. D2 or B) D0 vs. D14 comparisons from the DIA data.**

**Figure S21: KEGG enrichment analysis of upregulated proteins in proteomes in A) D0 vs. D2 or B) D0 vs. D14 comparisons from the DIA data.**

**Figure S22: GO enrichment analysis of proteomes from individual experimental groups in DIA data.** GO enrichment is classified by A) Biological Process, and B) Cellular Component.

**Figure S23: Molecular function GO enrichment analysis of proteomes from individual experimental groups in the DIA data.**

**Figure S24: GO enrichment of proteomes from D0 vs. D2 or D14 comparisons in DIA data.** GO enrichment is classified by A) Biological Process, B) Cellular Component, and C) Molecular Function as depicted in the plots.

**Figure S25: GO enrichment analysis of upregulated proteins from proteomes in D0 vs. D2 or D14 comparisons from the DIA data.** GO enrichment is classified by A) Biological Process, B) Cellular Component, and C) Molecular Function as depicted in the plots.

**Figure S26: GO enrichment analysis of downregulated proteins from proteomes in D0 vs. D2 or D14 comparisons from the DIA data.** GO enrichment is classified by A) Biological Process, B) Cellular Component, and C) Molecular Function as depicted in the plots.

**Figure S27: Spearman’s correlation coefficient (ρ) and data distribution plots among replicates for days, 2, 4, 6, 8, 10, 11, 12, and 14 in PRM data.**

**Figure S28: A) Distribution of MS2 data points across the chromatographic peak and B) correlation matrix among experimental groups from the PRM data.**

**Figure S29: Number of A) peptides and B) proteins, and C) distribution of number of peptides per protein in PRM data.** Data represent the median ± SD of two biological replicates.

**Figure S30: Protein ranking for erythroid samples collected on days 0, 2, 4, 6, 8, 10, 11, 12, and 14 in PRM data.** The name of every protein in the ranking is depicted in each plot. The Y-axis represents the log2 number of protein copies per nucleus in the proteomes of each experimental group, whereas the X-axis depicts the abundance ranking for proteins in a given proteome.

**Figure S31: Trend line plot for protein members of cBAF complex based on the PRM data.** Trend line plot for ACTL6 (ACTL6A, and ACTL6B), BCL7 (BCL7A, BCL7B, and BCL7C), DPF (DPF1-3), SMARCA (SMARCA2, and SMARCA4), SMARCC (SMARCC1, and SMARCC2), SS18 (SS18 AND SS18L1), and SMARCE1 proteins. Data represent the median ± SD of two biological replicates.

## Supplementary Tables

**Table S1: List of annotated tables generated for each QuickProt notebook.** A detailed description of the folder location, names of output spreadsheet tables, and their contents is provided.

**Table S2: DIA method for sample processing in Eclipse Orbitrap LC-MS/MS.**

**Table S3: PRM method for sample processing in Eclipse Orbitrap LC-MS/MS.**

**Table S4: Metadata table containing the names of samples (used for ease in analysis) and their respective LC-MS/MS raw file names.**

**Table S5: List of peptides and QconCATs used for PRM quantification.**

**Table S6: Molecular weight (MW) of QconCATs.**

**Table S7: Protein amount (pg) per nucleus in each experimental group.**

**Table S8: Median number of copies per nucleus of protein members of the cBAF complex.** The median copies per nucleus were calculated first among QconCATs within each replicate, then these values were averaged across two biological replicates to calculate the final copies per nucleus. The minimum and maximum values for each protein across the time course are also represented in the table.

## Notes

### Competing Interest Statement

The authors have declared no competing interest.

