## Supplementary Figures for "QuickProt: A bioinformatics and visualization tool for DIA and PRM mass spectrometry-based proteomics datasets"

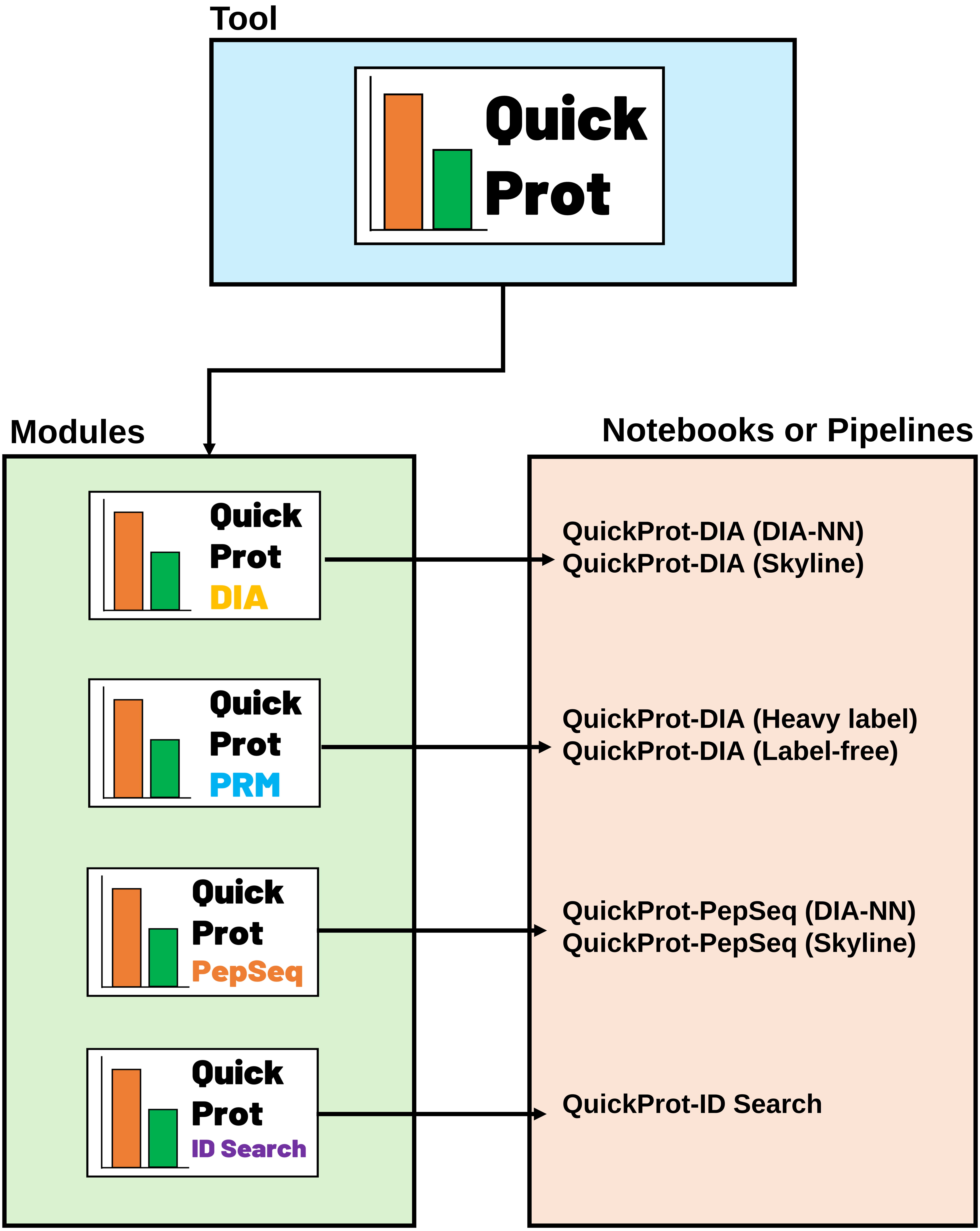


**Figure S1: Scheme of the QuickProt tool for proteomic data mining and visualization. QuickProt comprises five modules: QuickProt-DIA, QuickProt-PRM, QuickProt-PepSeq, and QuickProt-ID Search.** QuickProt-DIA consists of two notebooks or pipelines: QuickProt-DIA (DIA-NN) and QuickProt-DIA (Skyline). Additionally, QuickProt-PRM includes two notebooks: QuickProt-PRM (Heavy label) and QuickProt-PRM (Label-free). QuickProt-PepSeq features two pipelines: QuickProt-PepSeq (DIA-NN) and QuickProt-PepSeq (Skyline). Lastly, QuickProt-ID Search has a notebook of the same name.

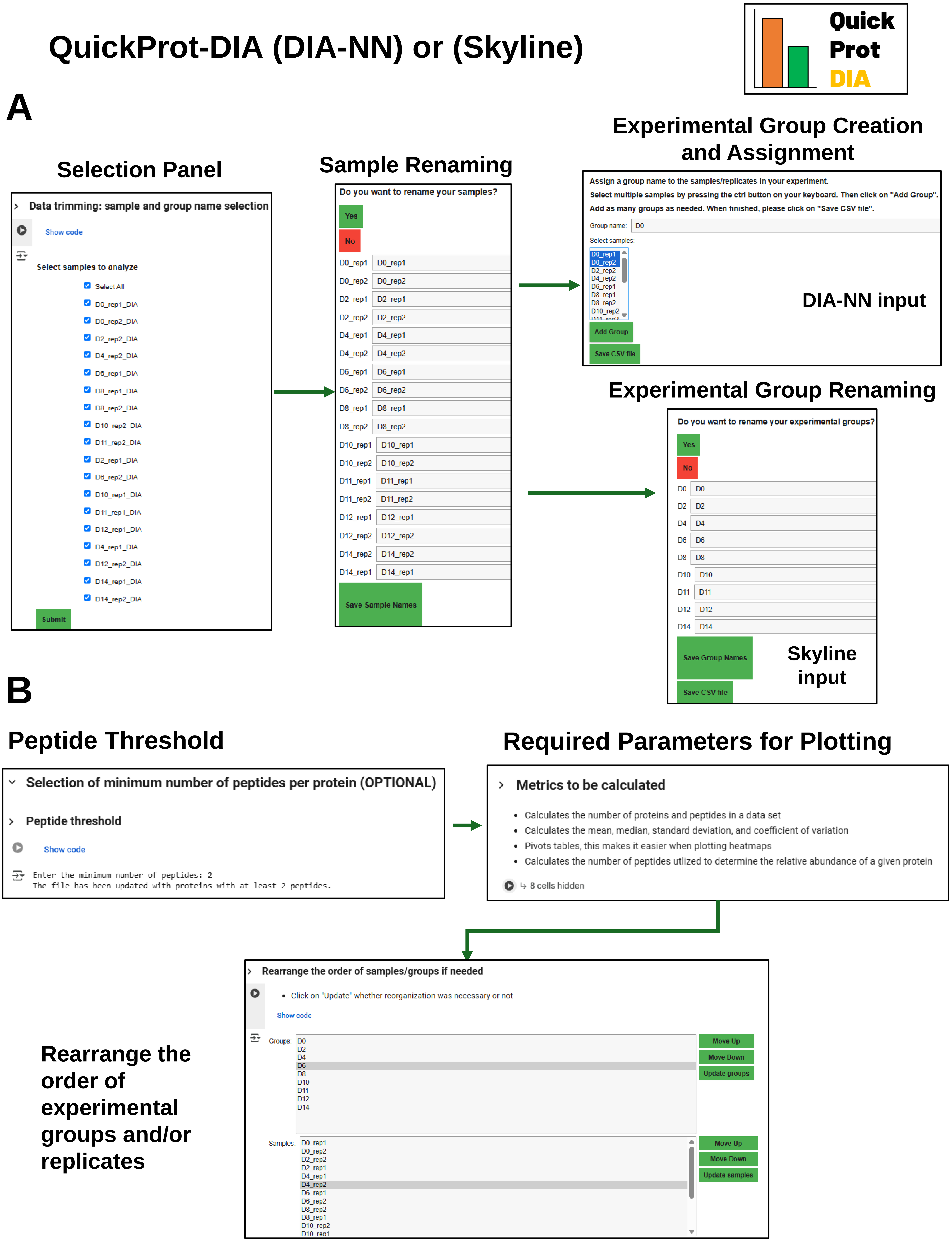


**Figure S2: Overview of the QuickProt-DIA (DIA-NN) or (Skyline) notebook data preprocessing interface.**

### QuickProt-DIA (DIA-NN) or (Skyline)

#### Quality Control

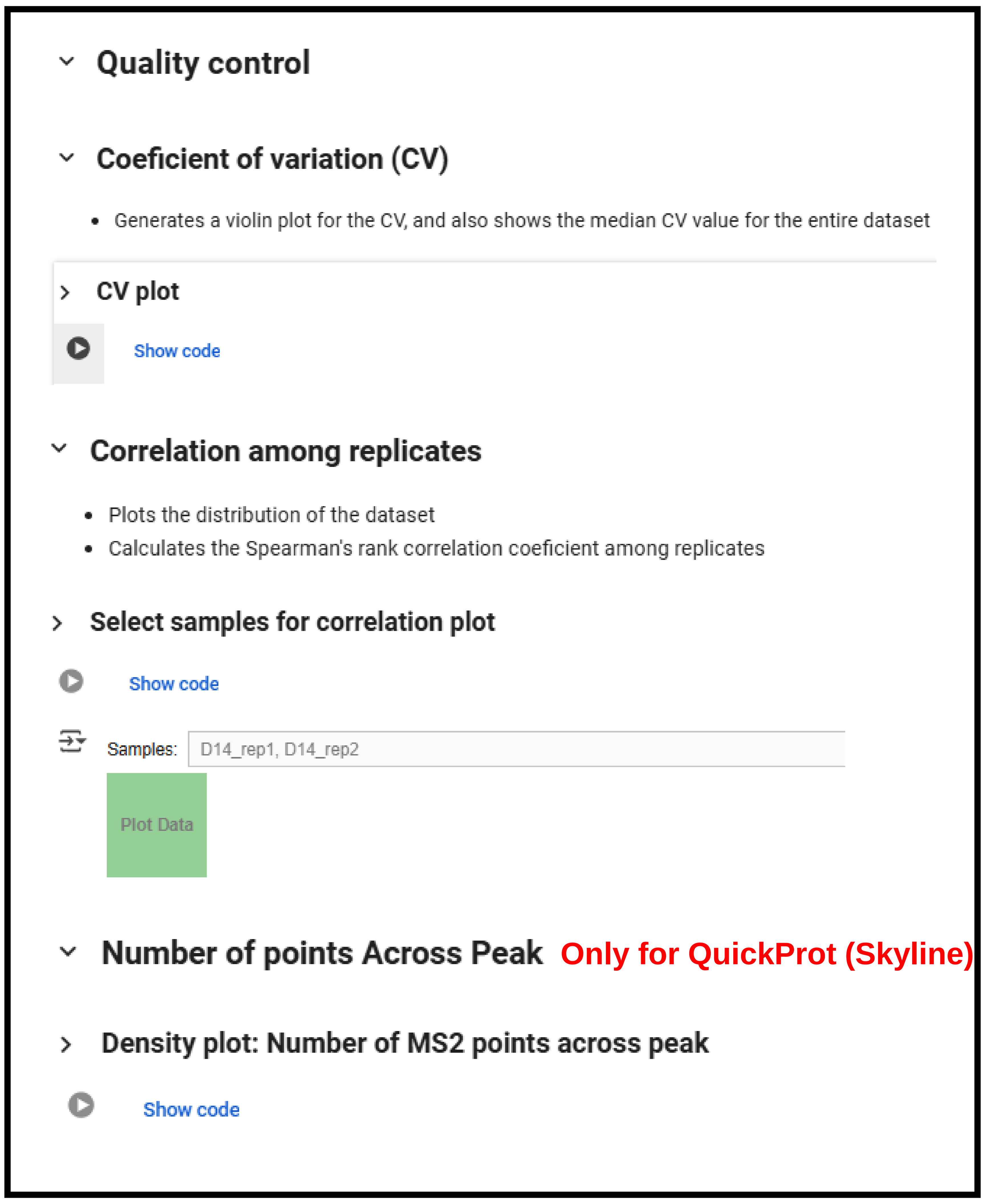


#### Peptide and Protein Yields

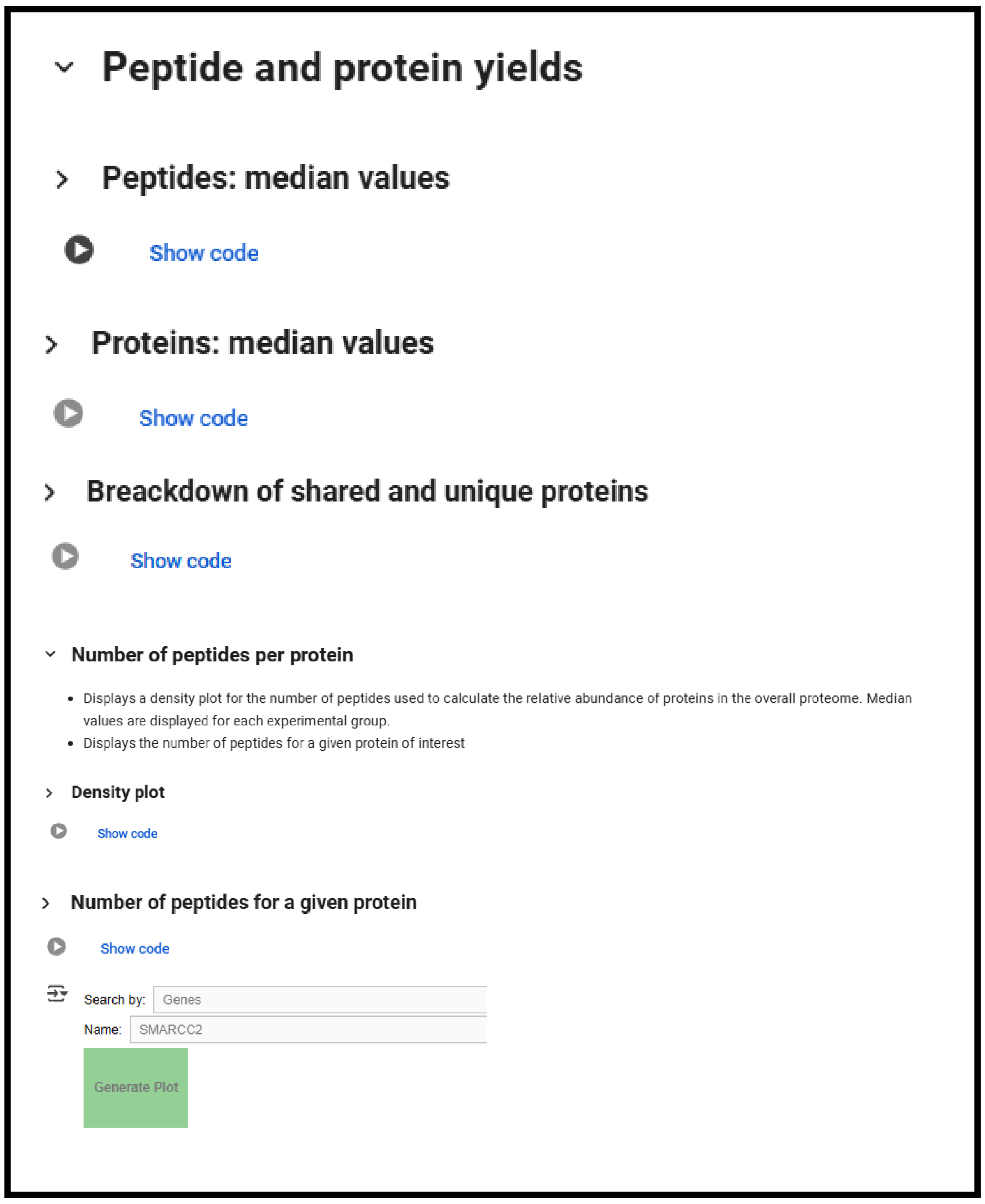


#### Exploratory Analysis

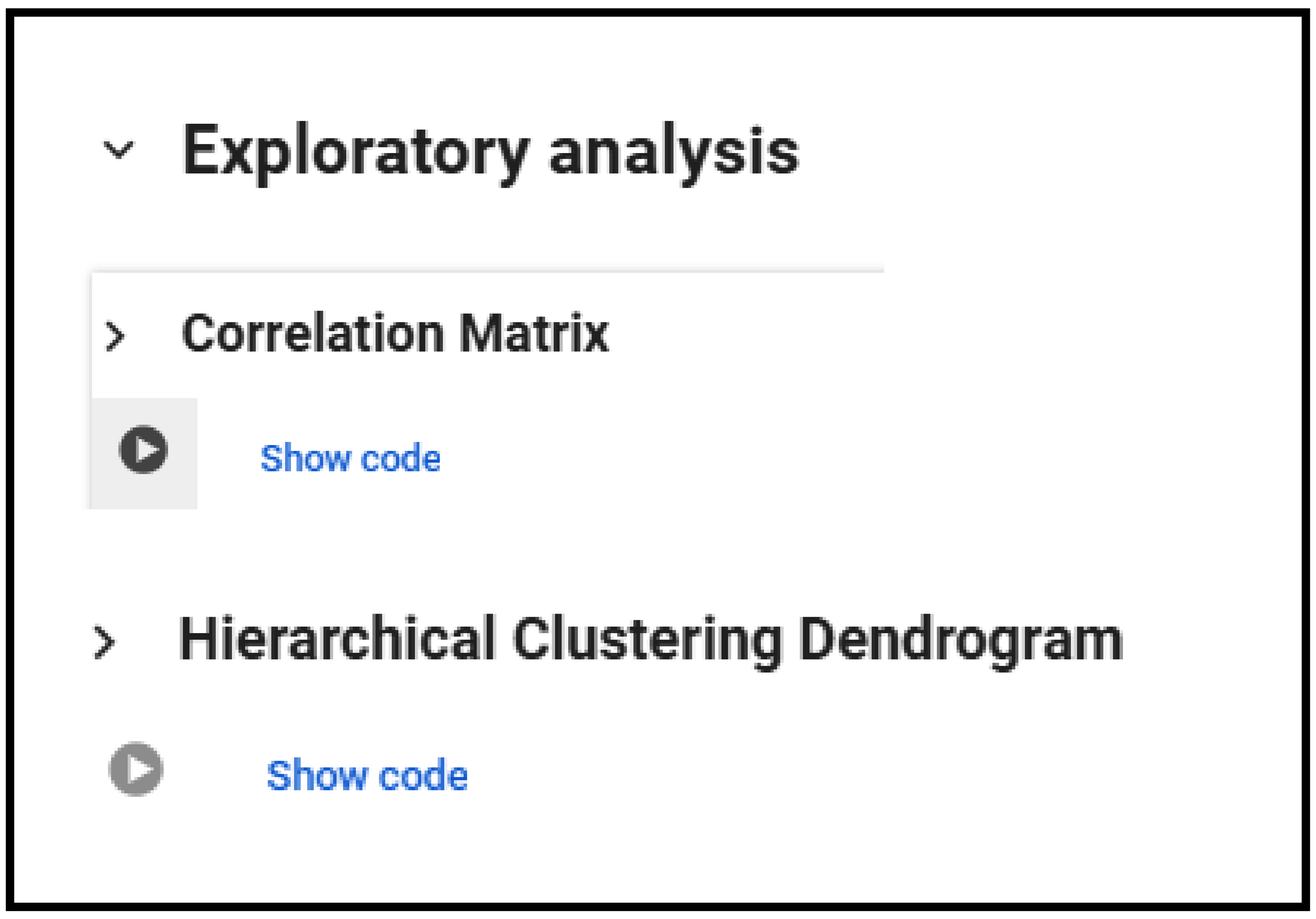


**Figure S3: Overview of the quality control, peptide and protein yields, and exploratory analysis interface in the QuickProt-DIA (DIA-NN) or (Skyline) notebooks.**

### QuickProt-DIA (DIA-NN) or (Skyline)

#### Protein Abundance

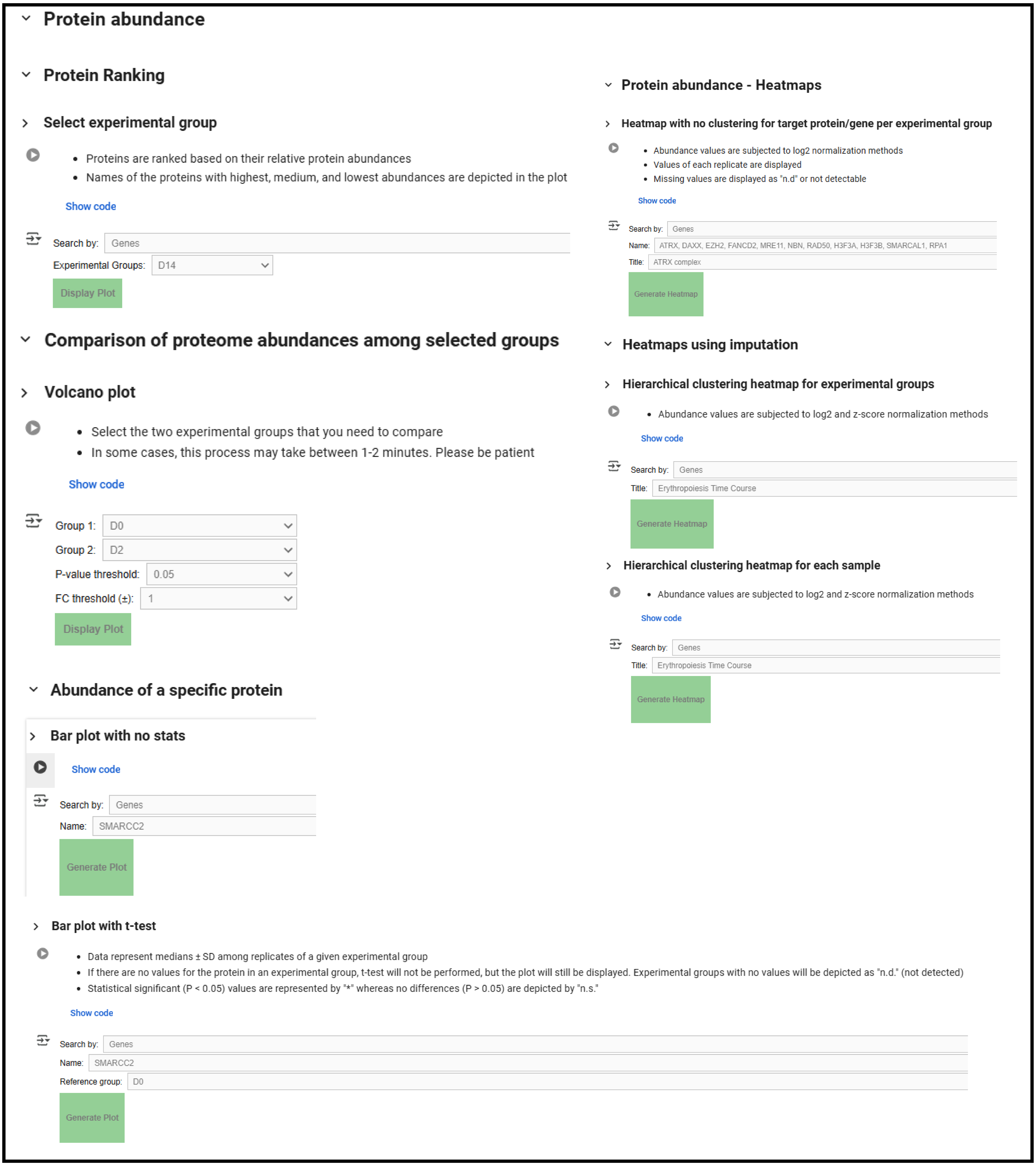


**Figure S4: Overview of the protein abundance analysis interface in the QuickProt-DIA (DIA-NN) or (Skyline) notebooks.**

### QuickProt-DIA (DIA-NN) or (Skyline)

#### Enrichment Analysis

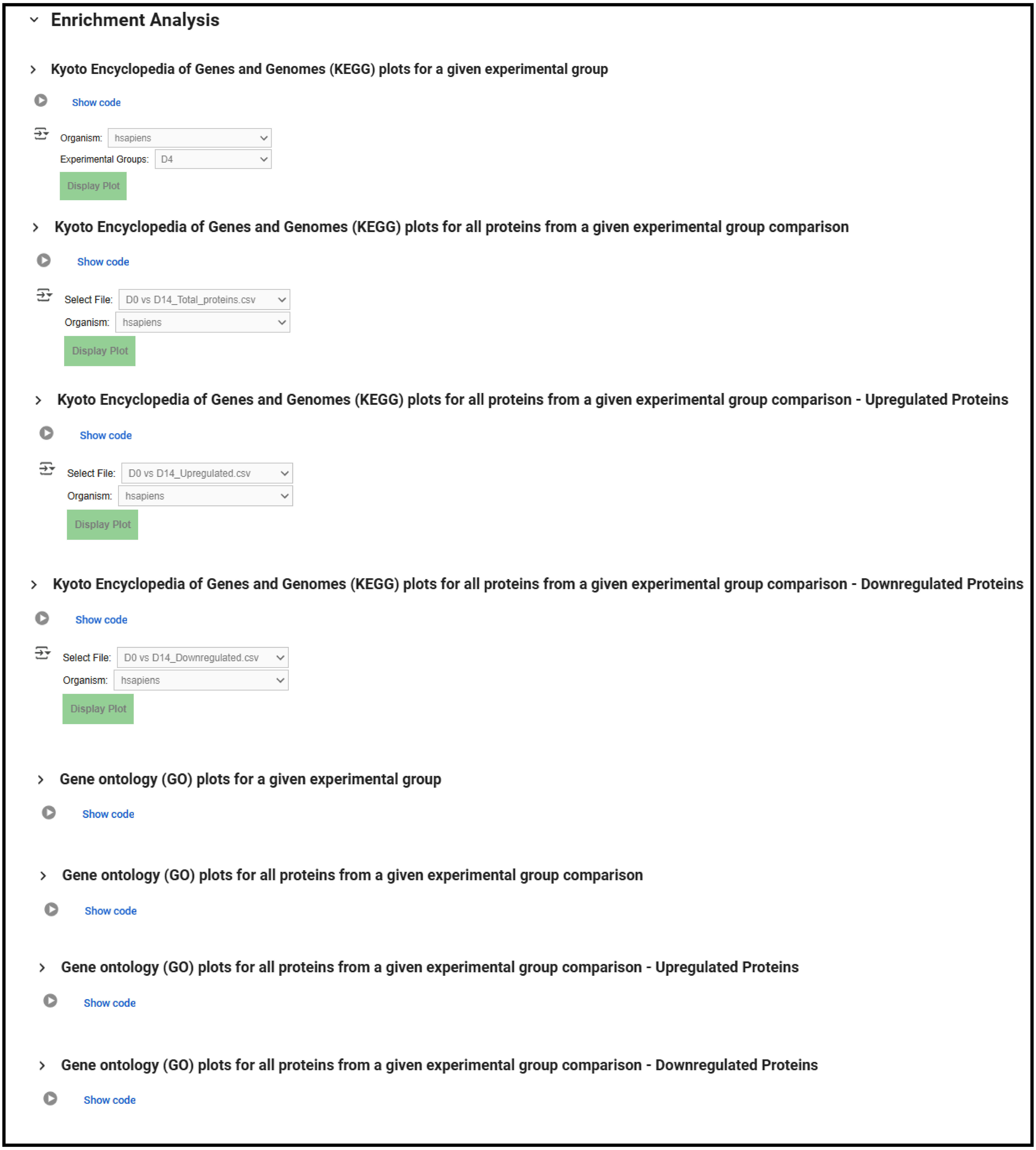


**Figure S5: Overview of the enrichment analysis interface in the QuickProt-DIA (DIA-NN) or (Skyline) notebooks.**

### Preprocessing in QuickProt-PRM (Label-free)

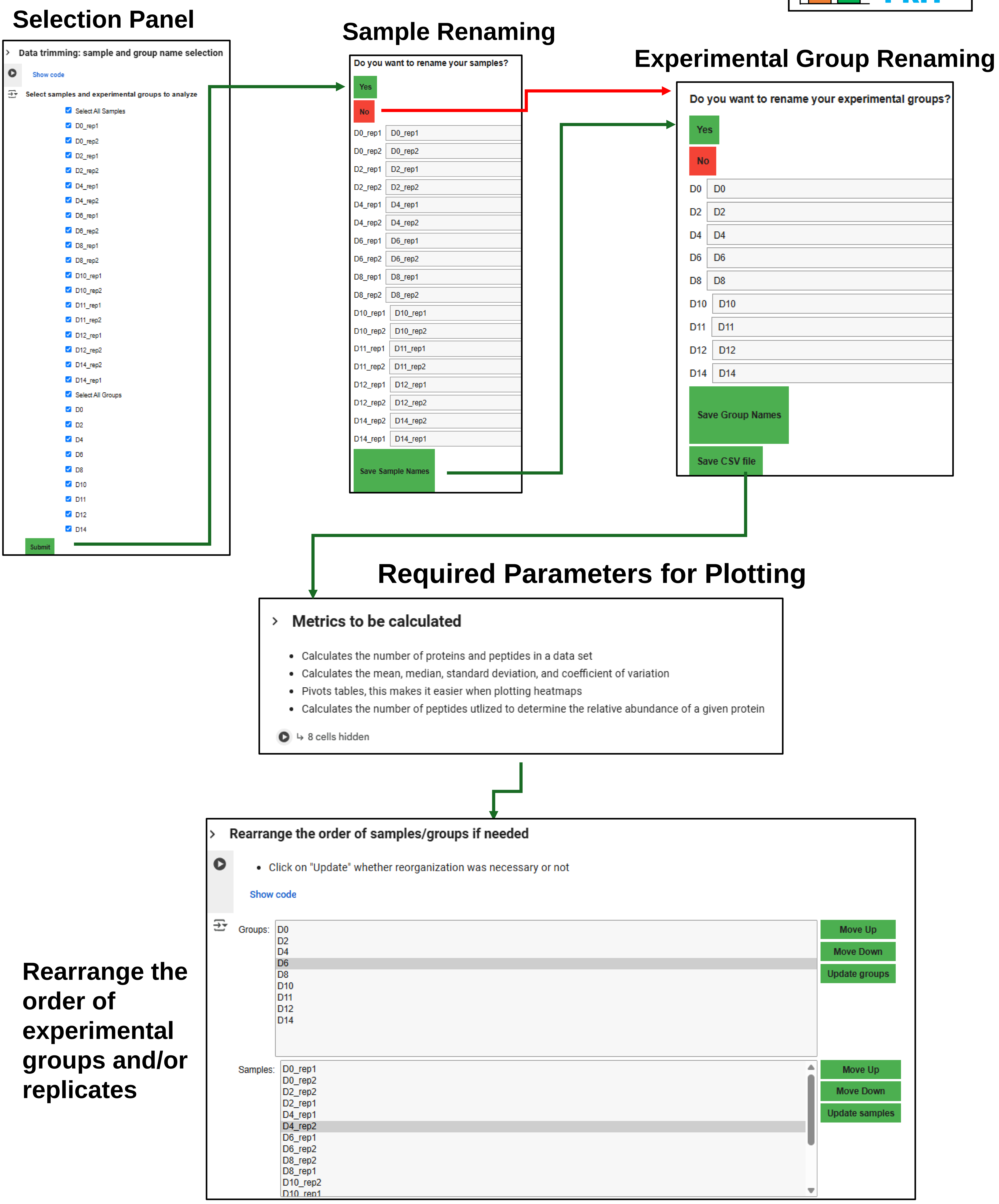


**Figure S6: Overview of the data preprocessing interface in the QuickProt-PRM (Label-free) notebook.**

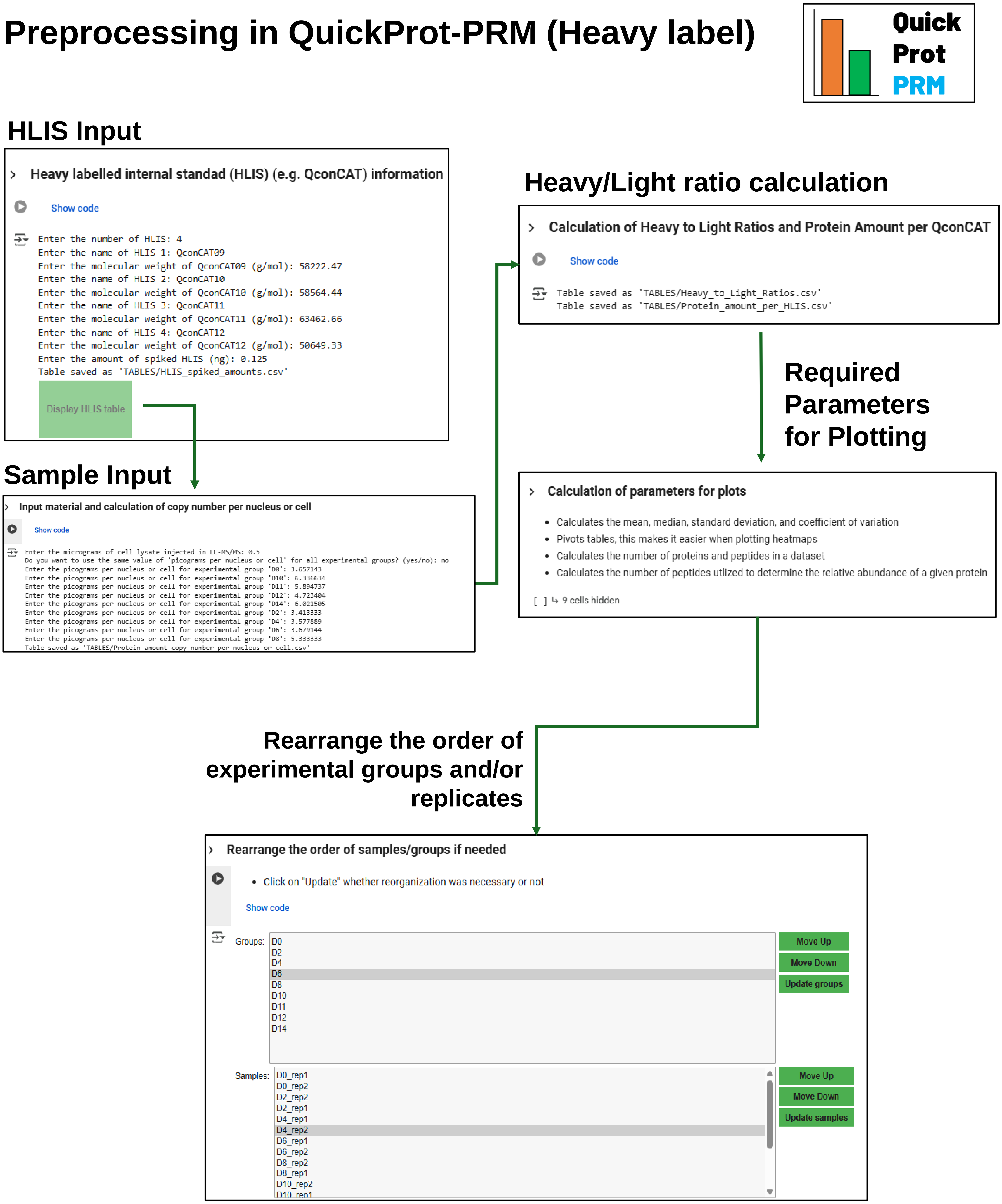


**Figure S7: Overview of the data preprocessing interface in the QuickProt-PRM (Heavy label) notebook.**

### QuickProt-PRM (Label-free) or (Heavy label)

#### Quality Control

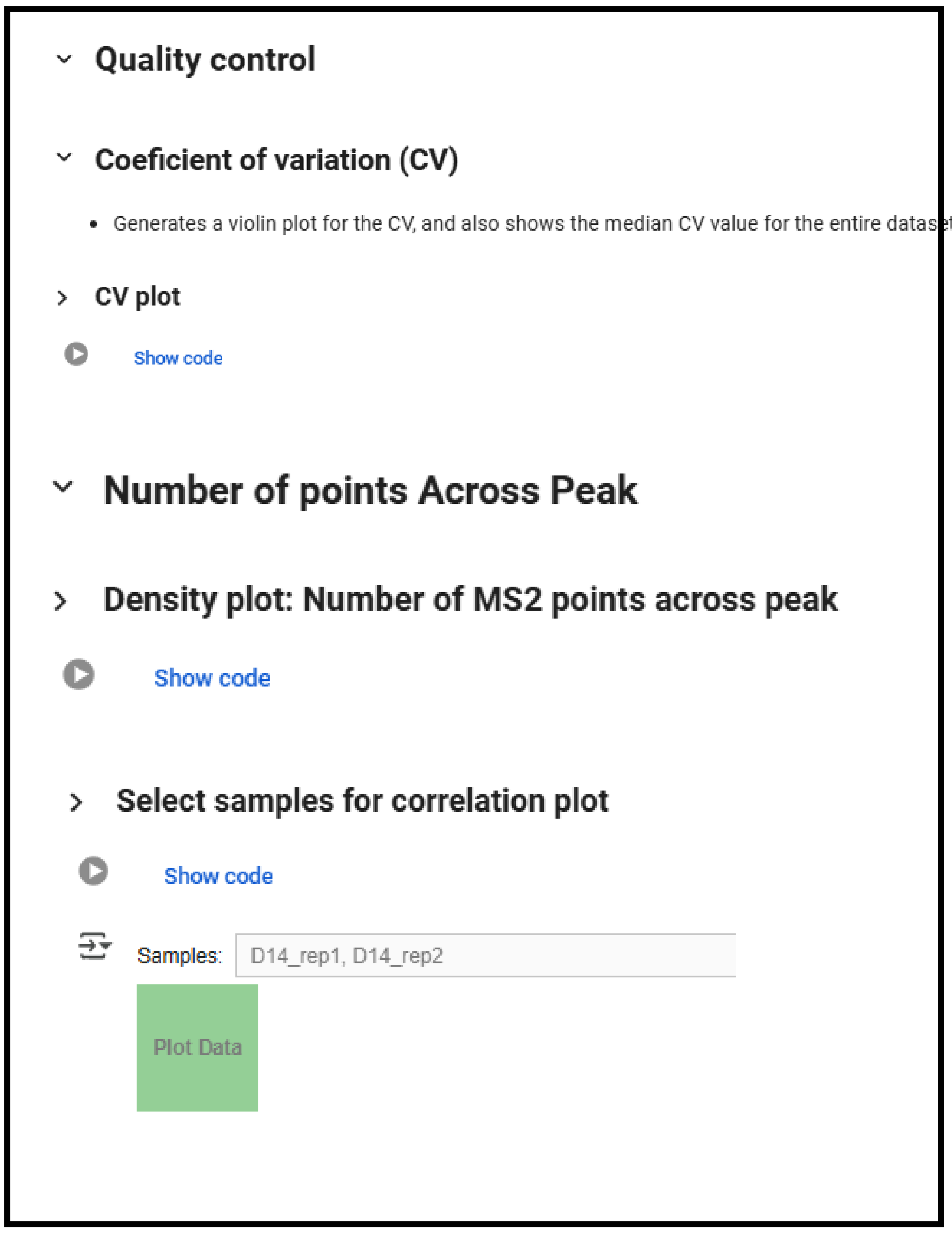


#### Peptide and Protein Yields

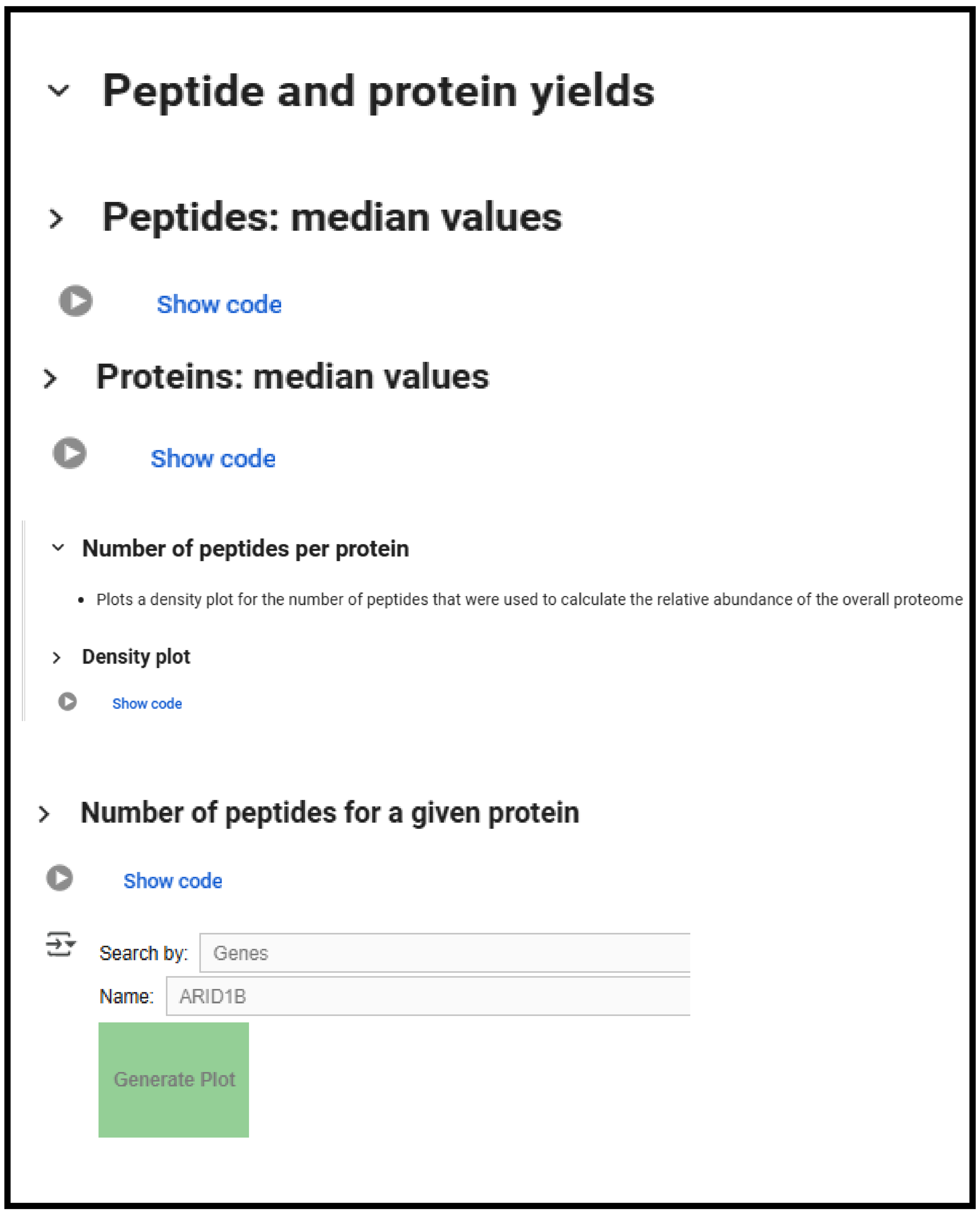


#### Exploratory Analysis

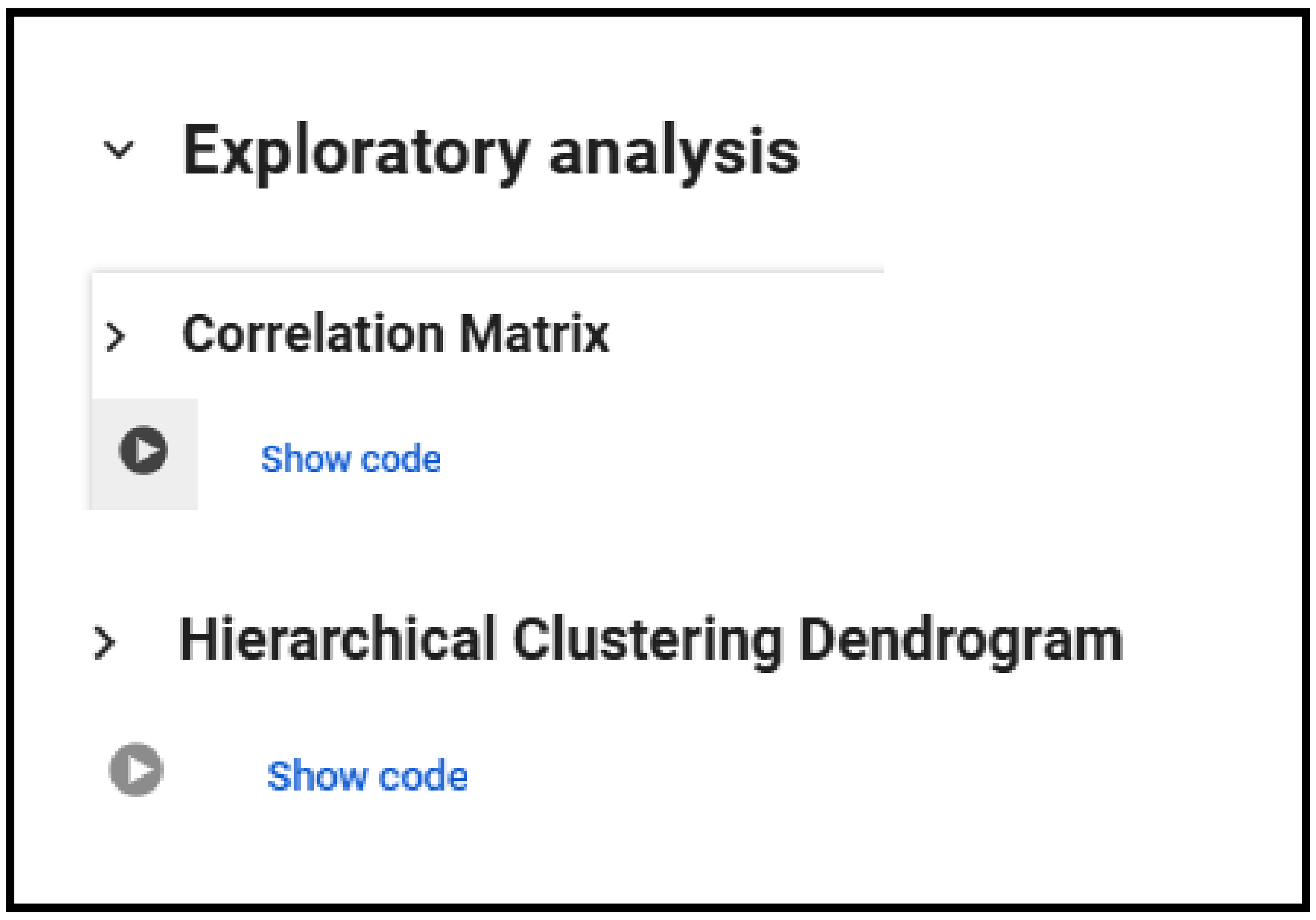


**Figure S8: Overview of the quality control, peptide and protein yields, and exploratory analysis interface in the QuickProt-PRM (Label-free) or (Heavy label) notebooks.**

### QuickProt-PRM (Label-free)

#### Protein Abundance

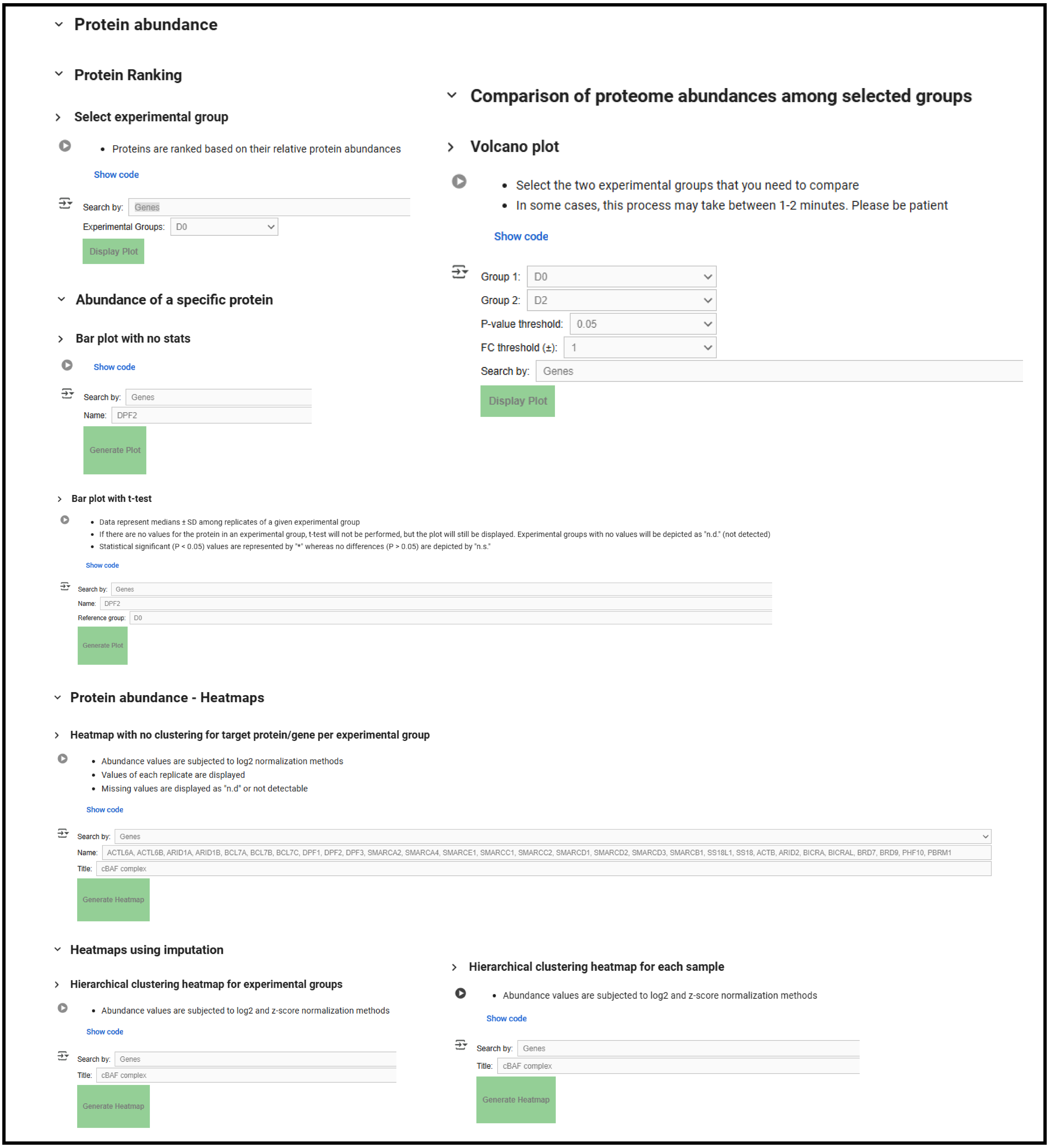


**Figure S9: Overview of the protein abundance analysis interface in the QuickProt-PRM (Label-free) notebooks.**

### QuickProt-PRM (Heavy label)

#### Protein Abundance

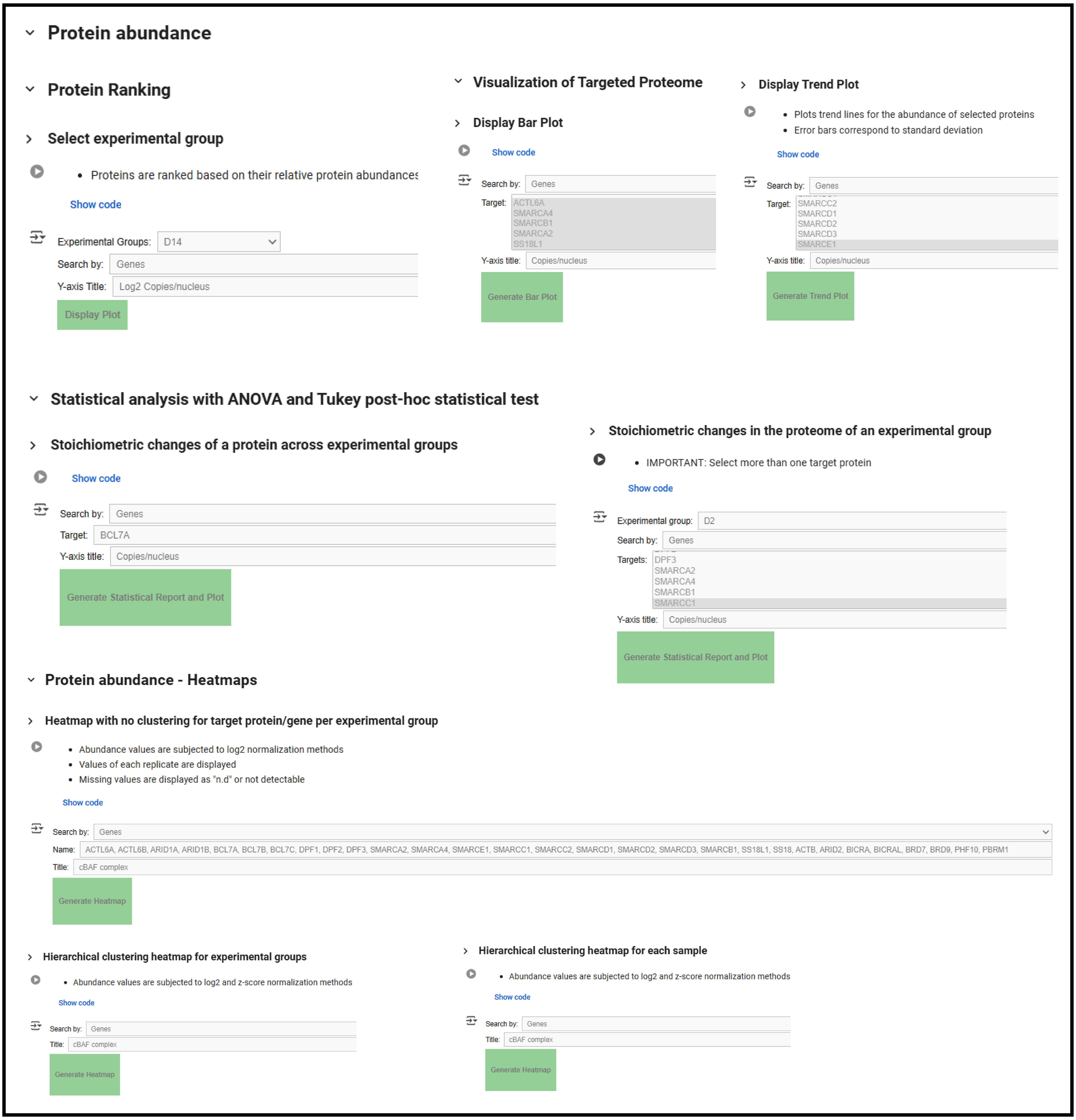


**Figure S10: Overview of the protein abundance analysis interface in the QuickProt-PRM (Heavy label) notebooks.**

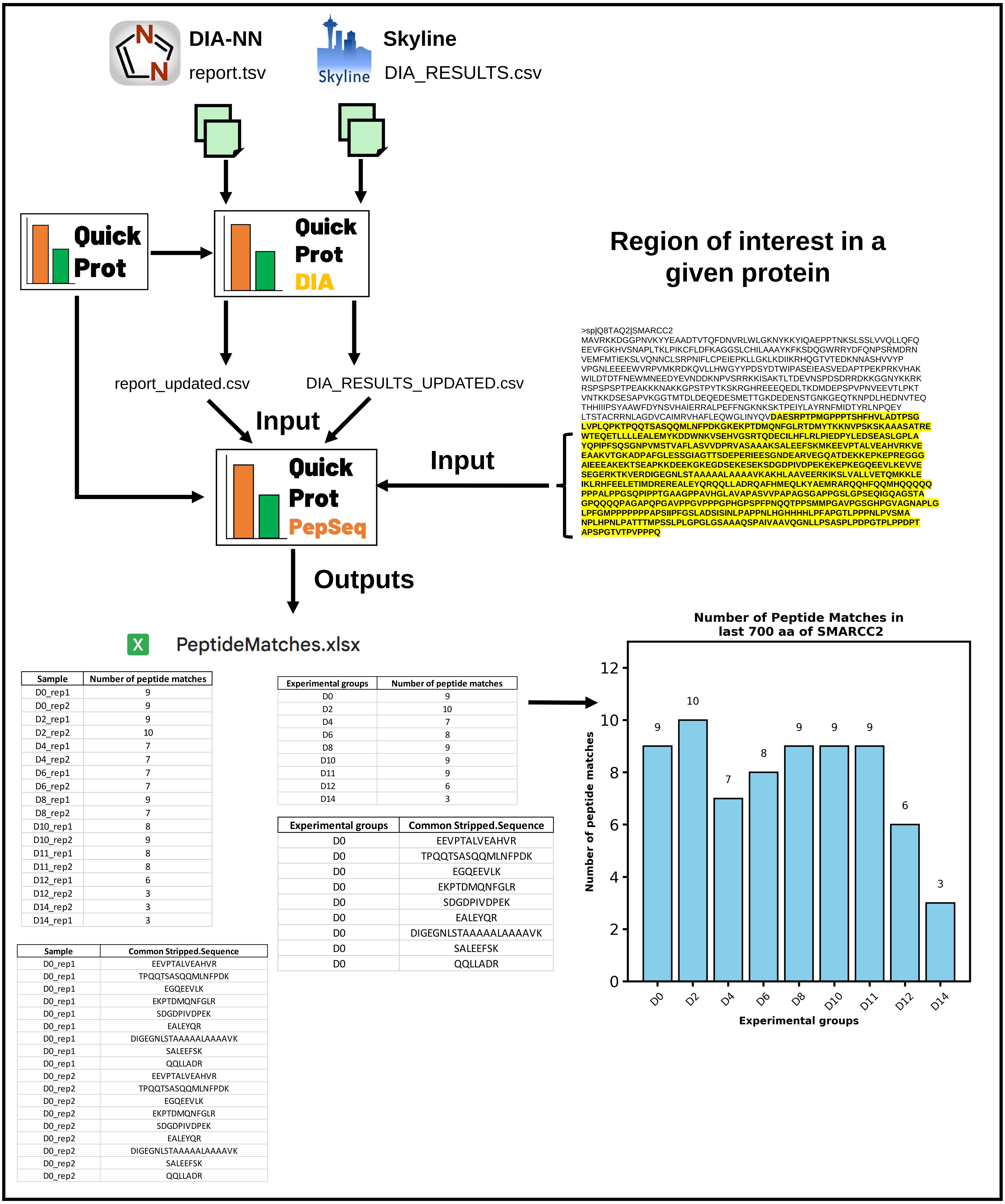


**Figure S11: Overview of QuickProt-PepSeq workflow.** Outputs from DIA-NN or Skyline, coupled with QuickProt-DIA, can be imported into the QuickProt-PepSeq notebook (part of the QuickProt notebook series). A given region of interest (e.g., a protein domain) is then entered into the notebook to be mapped against the DIA data of a specific experiment. As a result, an Excel file, 'PeptideMatches.xlsx' will be generated. This file contains, in different tabs, the number of peptide matches per sample and per experimental group, as well as their respective amino acid sequences. The notebook also includes an option for plotting a bar graph showing the number of peptide matches in each experimental group.

Target_gene_list.csv

| Gene name |
|---|
| ACTB |
| ACTL6A |
| ACTL6B |
| ARID1A |
| ARID1B |
| ARID2 |
| BCL7A |
| BCL7B |
| BCL7C |
| BICRA |
| BICRAL |
| BRD7 |
| BRD9 |
| DPF1 |
| DPF2 |
| DPF3 |
| PBRM1 |
| PHF10 |
| SMARCA2 |
| SMARCA4 |
| SMARCB1 |
| SMARCC1 |
| SMARCC2 |
| SMARCD1 |
| SMARCD2 |
| SMARCD3 |
| SMARCE1 |
| SS18 |
| SS18L1 |

Quick Prot

Quick Prot ID Search

Target_gene_ProteinID_list.csv

| Gene name | Protein IDs |
|---|---|
| ACTB | P60709 |
| ACTL6A | O96019 |
| ACTL6B | O94805 |
| ARID1A | O14497 |
| ARID1B | Q8NFD5 |
| ARID2 | Q68CP9 |
| BCL7A | Q4VC05 |
| BCL7B | Q9BQE9 |
| BCL7C | Q8WUZ0 |
| BICRA | Q9NZM4 |
| BICRAL | Q6AI39 |
| BRD7 | Q9NPI1 |
| BRD9 | Q9H8M2 |
| DPF1 | Q92782 |
| DPF2 | Q92785 |
| DPF3 | Q92784 |
| PBRM1 | Q86U86 |
| PHF10 | Q8WUB8 |
| SMARCA2 | P51531 |
| SMARCA4 | P51532 |
| SMARCB1 | Q12824 |
| SMARCC1 | Q92922 |
| SMARCC2 | Q8TAQ2 |
| SMARCD1 | Q96GM5 |
| SMARCD2 | Q92925 |
| SMARCD3 | Q6STE5 |
| SMARCE1 | Q969G3 |
| SS18 | Q15532 |
| SS18L1 | O75177 |

**Figure S12: Overview of QuickProt-ID Search workflow.** The user inputs a list of genes of interest into QuickProt-ID. Using the Unipressed (Uniprot REST) library, the notebook has been adapted to generate a CSV table with their respective protein IDs.

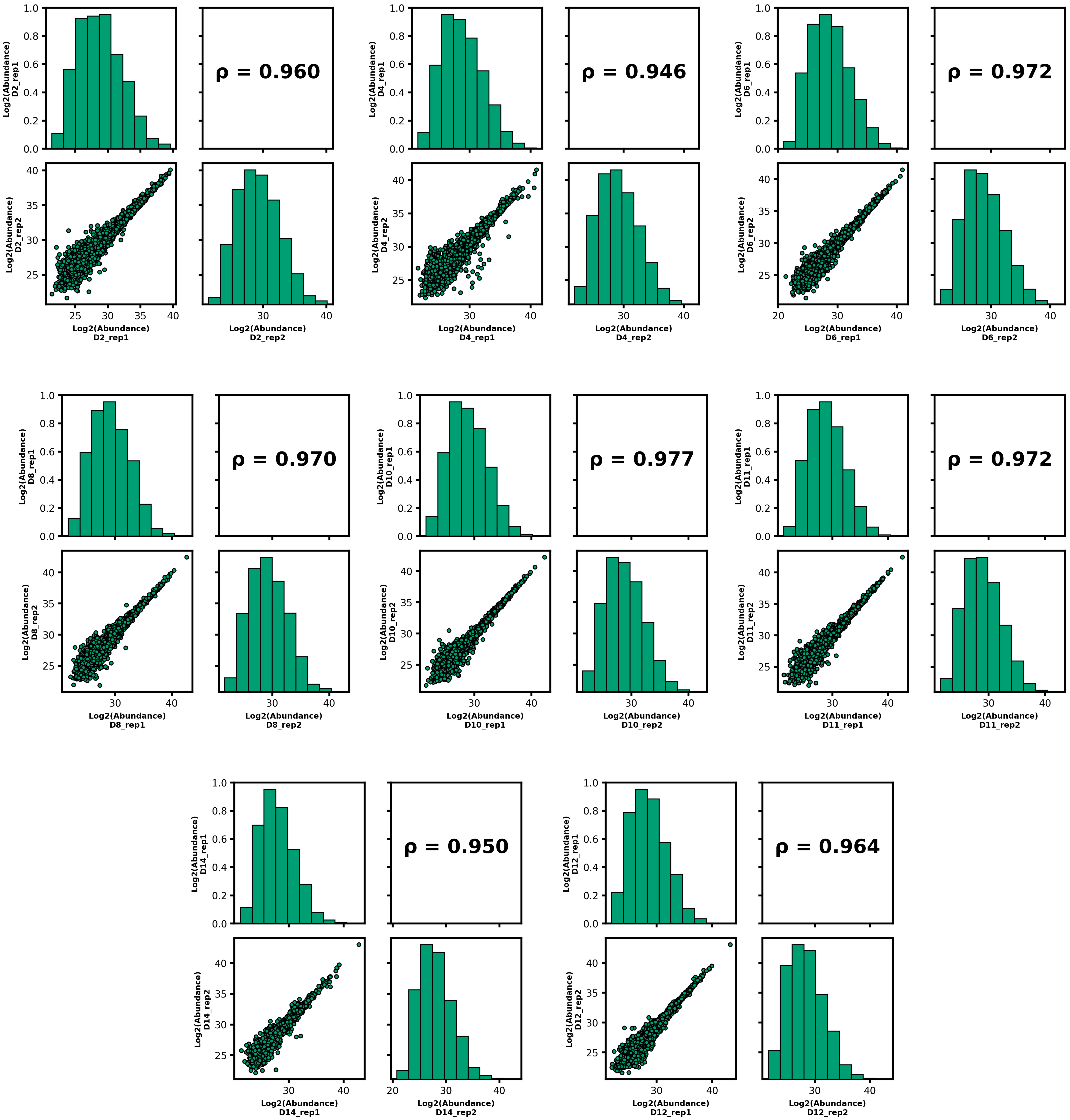


**Figure S13: Spearman’s correlation coefficient (ρ) and data distribution plots among replicates for days, 2, 4, 6, 8, 10, 11, 12, and 14 in DIA-data.**

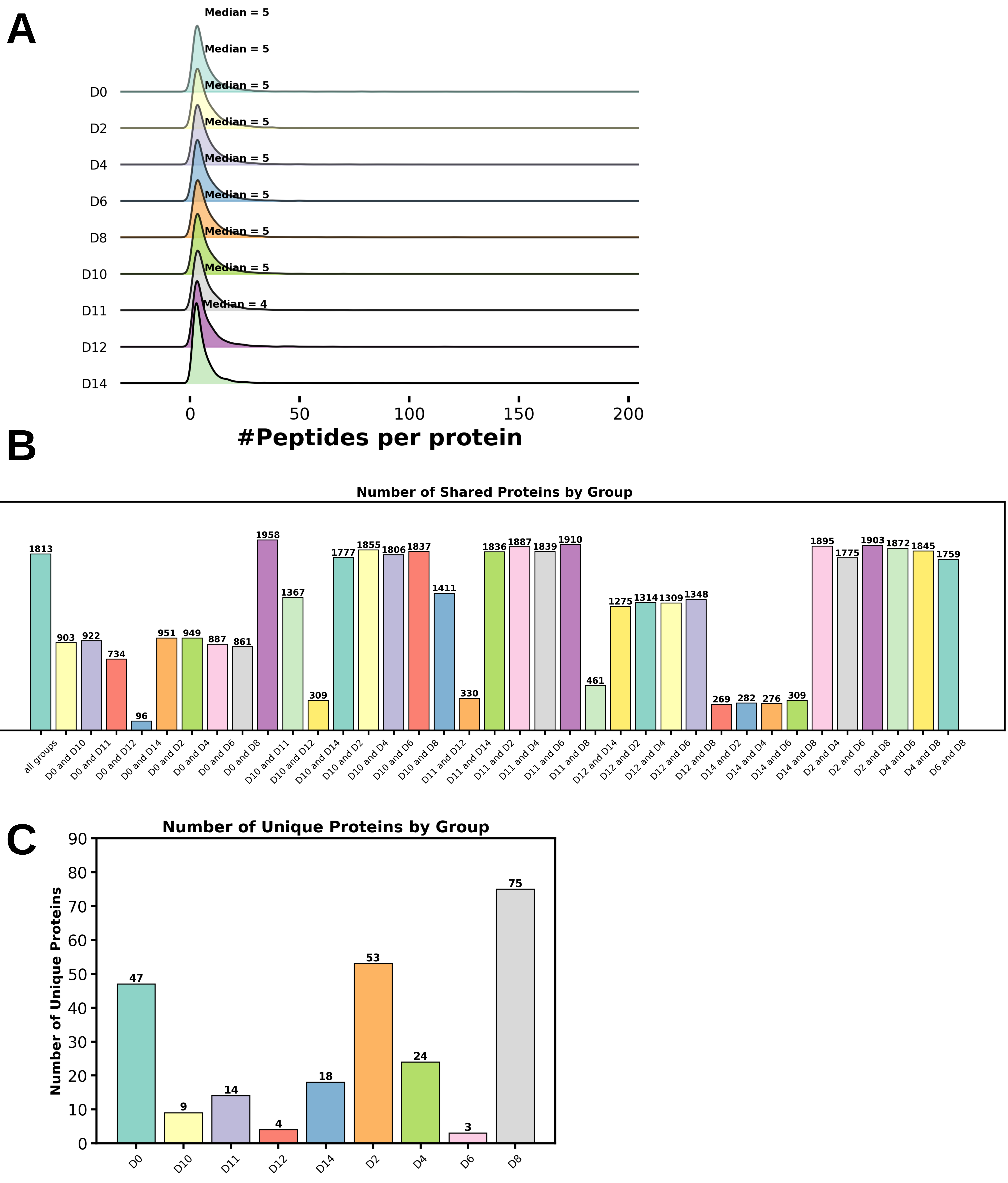


**Figure S14: Distribution of the number of peptides per protein and bar plots for shared and unique proteins in DIA data.** A) The number of peptides per protein plot depicts the distribution of values as well as the median value for each experimental group. Bar plots depict the median values for B) shared and C) unique proteins. A summary table with the names and numbers of proteins is generated and stored in a subfolder called 'SHARED_UNIQUE_PROTEINS' within the 'TABLES' folder. Additionally, within the same subfolder, another folder called 'Extracted' is generated to extract and store tables listing the proteins that are shared or unique for a given comparison or experimental group.

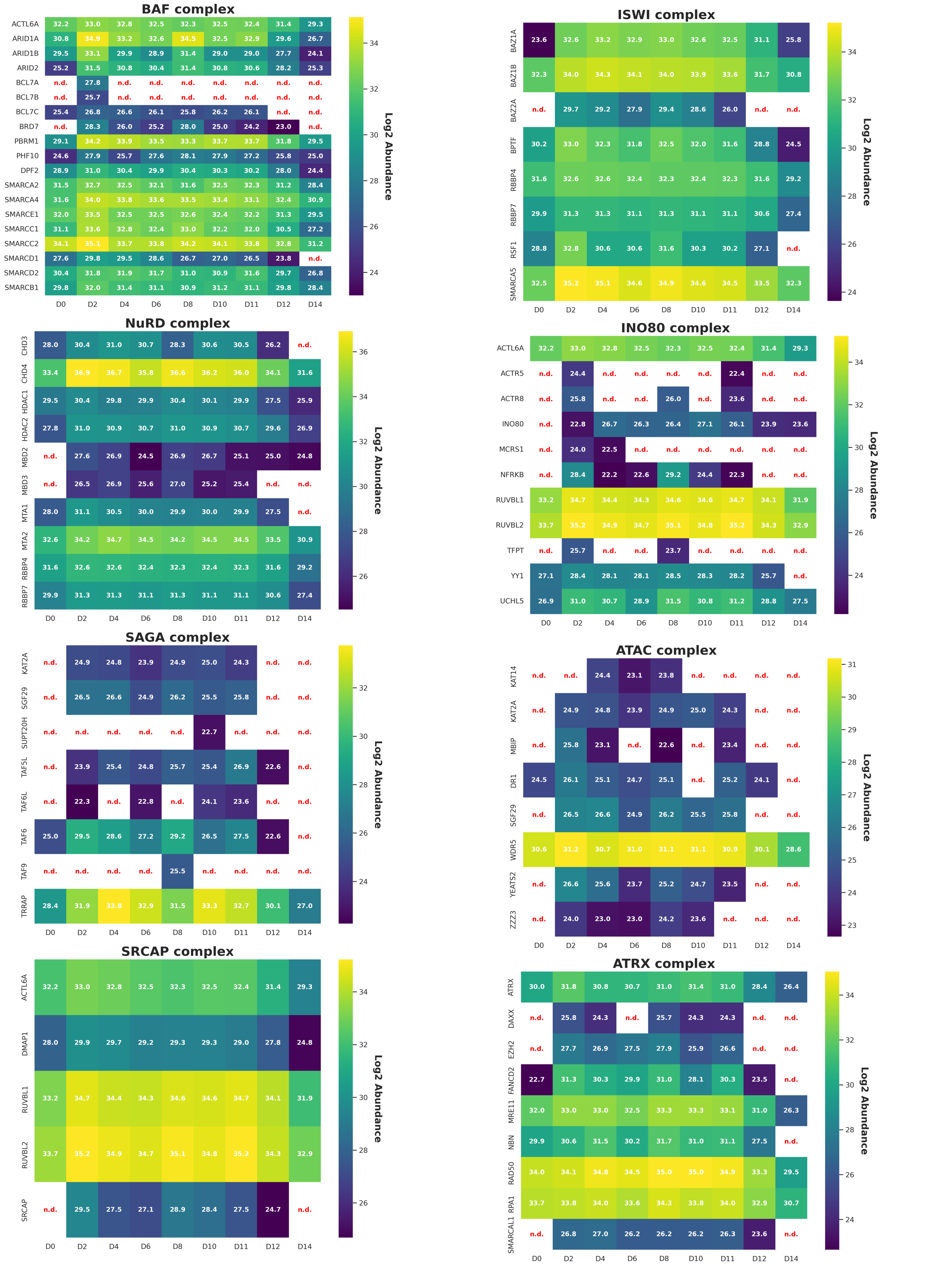


**Figure S15: Heatmaps show the relative abundances of proteins in the BAF, ISWI, NuRD, INO80, SAGA, ATAC, SRCAP, and ARTX chromatin remodeling complexes during the time course in the DIA data.** Log2 normalization was applied to the abundance values for proteins from days 0 to 14. The abbreviation ‘n.d.’ stands for ‘not detected’, indicating values that were not found in the datasets.

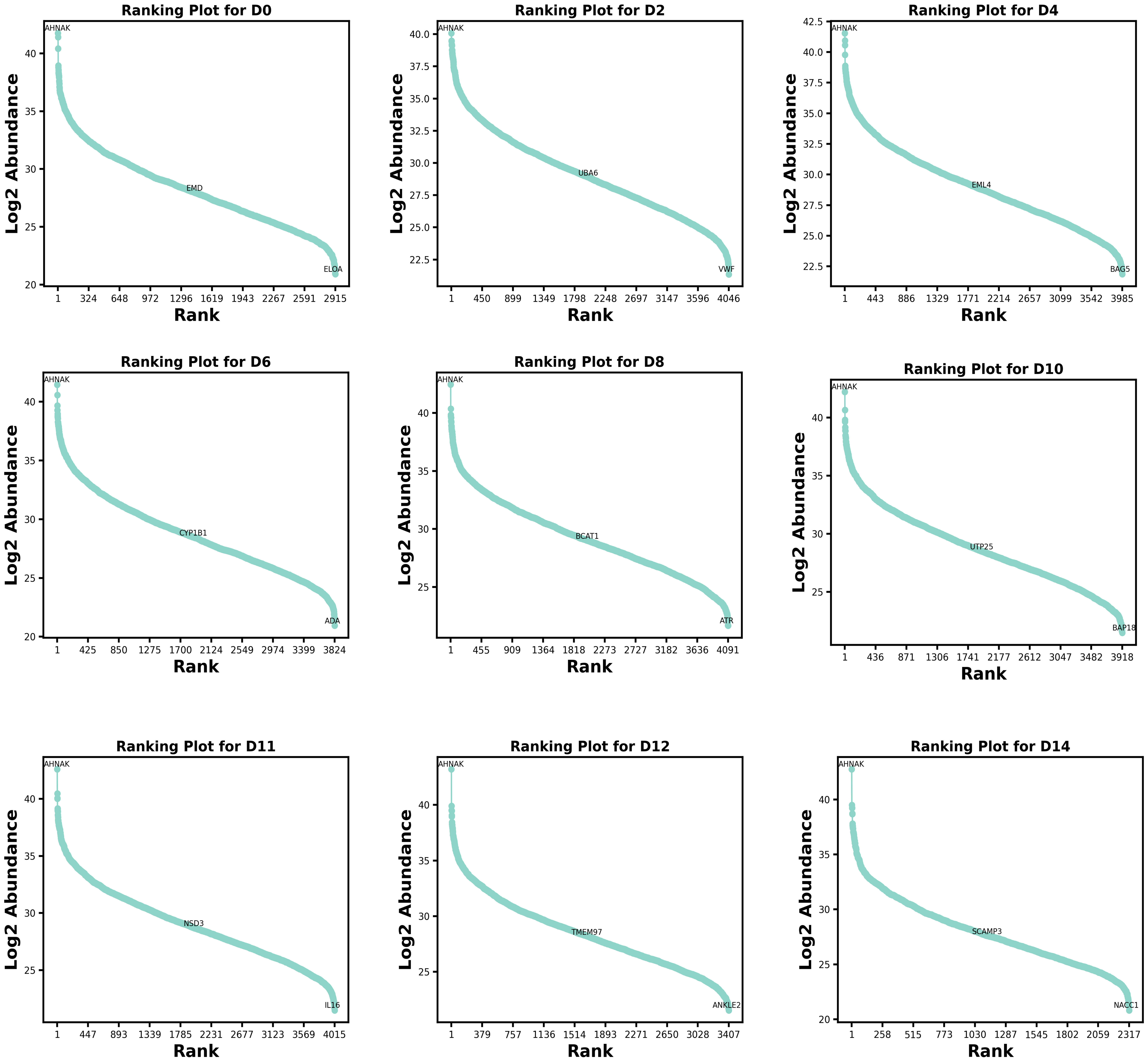


**Figure S16: Protein ranking for erythroid samples collected on days 0, 2, 4, 6, 8, 10, 11, 12, and 14 in DIA data.** The Y-axis represents the log2 abundance of proteins in the proteomes of each experimental group, whereas the X-axis depicts the abundance ranking for proteins in a given proteome. The names of the maximum, median, and minimum-ranking proteins are shown in the plot.

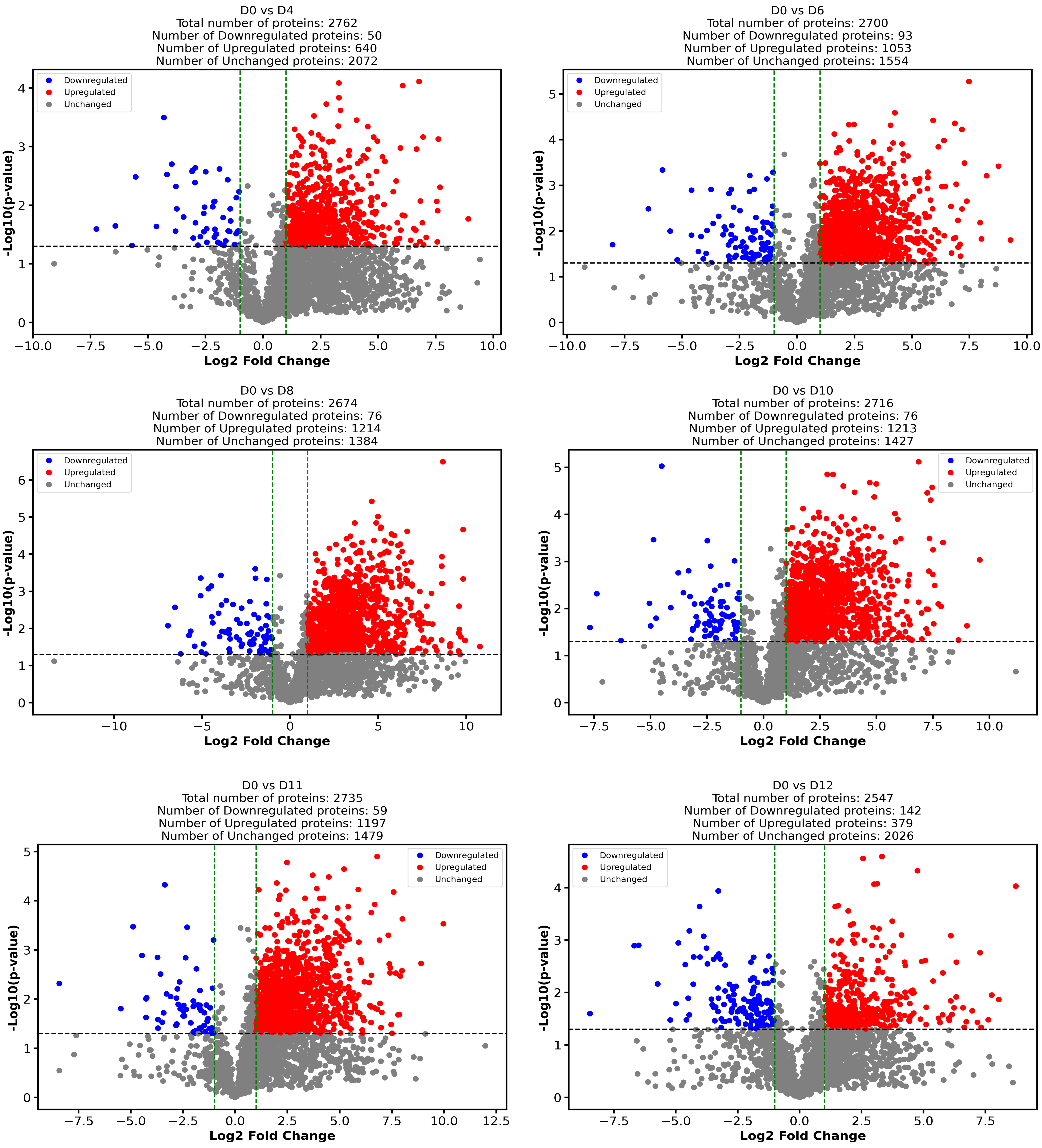


**Figure S17: Volcano plots for differential expression of D0 vs. D4, D6, D8, D10, D11, or D12 from the DIA data.** The volcano plots depict a *P*-value ≤ 0.05 and a fold-change (FC) threshold of 1.0. In the upper part of the plot, beside the title, a detailed description of the total number of upregulated, downregulated, and non-significantly changed proteins is shown.

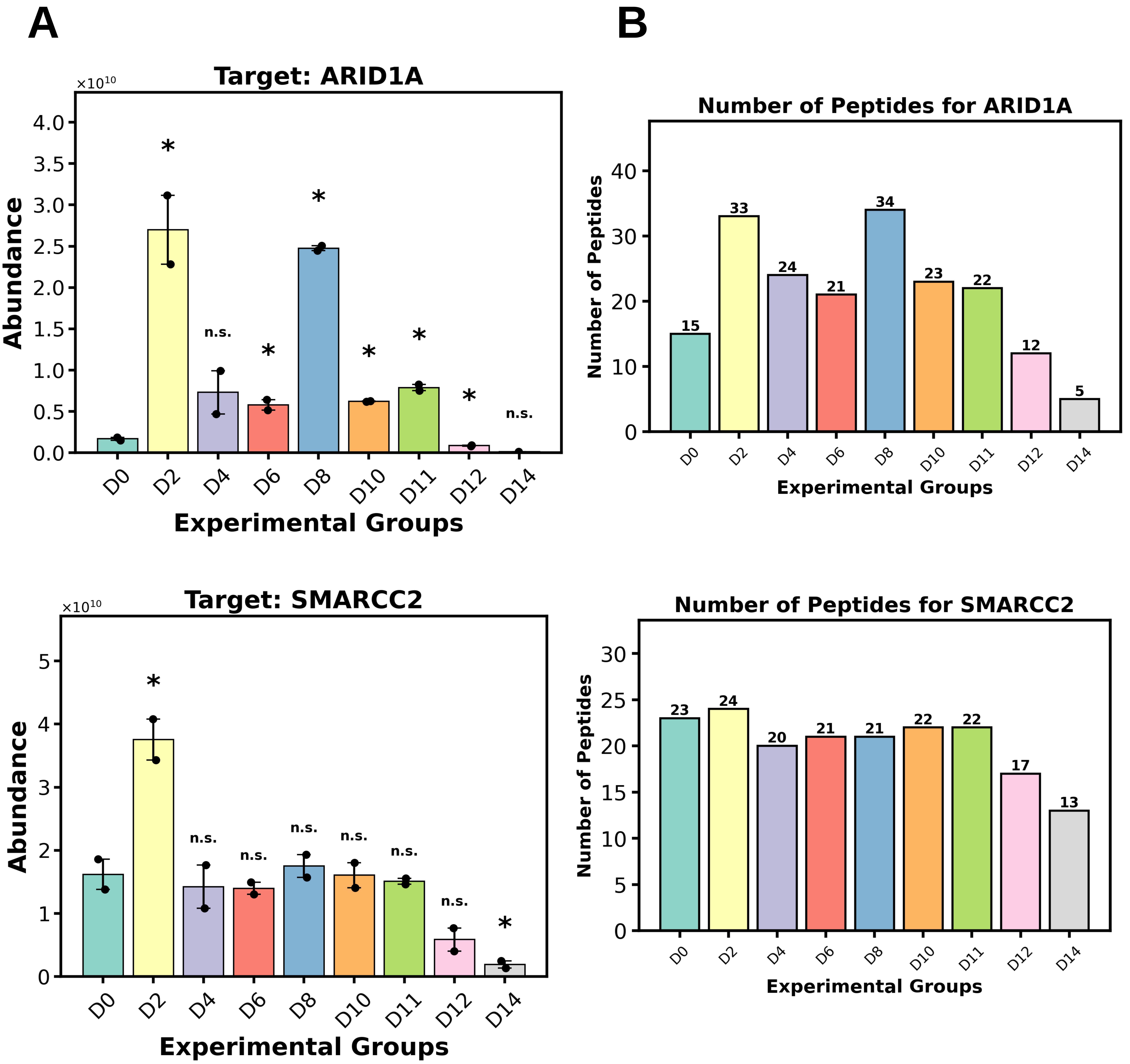


**Figure S18: Bar plots for protein abundance and peptide numbers for selected proteins in the DIA data of the erythroid samples.** A) The abundance of ARID1A (upper panel) and SMARCC2 (lower panel) proteins is displayed in bar plots. Statistical analysis by *t*-test was performed on the datasets, using Day 0 as the reference group for comparison. Asterisks represent statistically significant differences ($P \leq 0.05$), whereas 'n.s.' indicates non-significant differences. These statistical representations were added automatically by the code in the QuickProt-DIA notebook. B) Number of peptides supporting the abundance estimate for ARID1A and SMARCC2 in each experimental group. Data represent the median ± SD of two biological replicates.

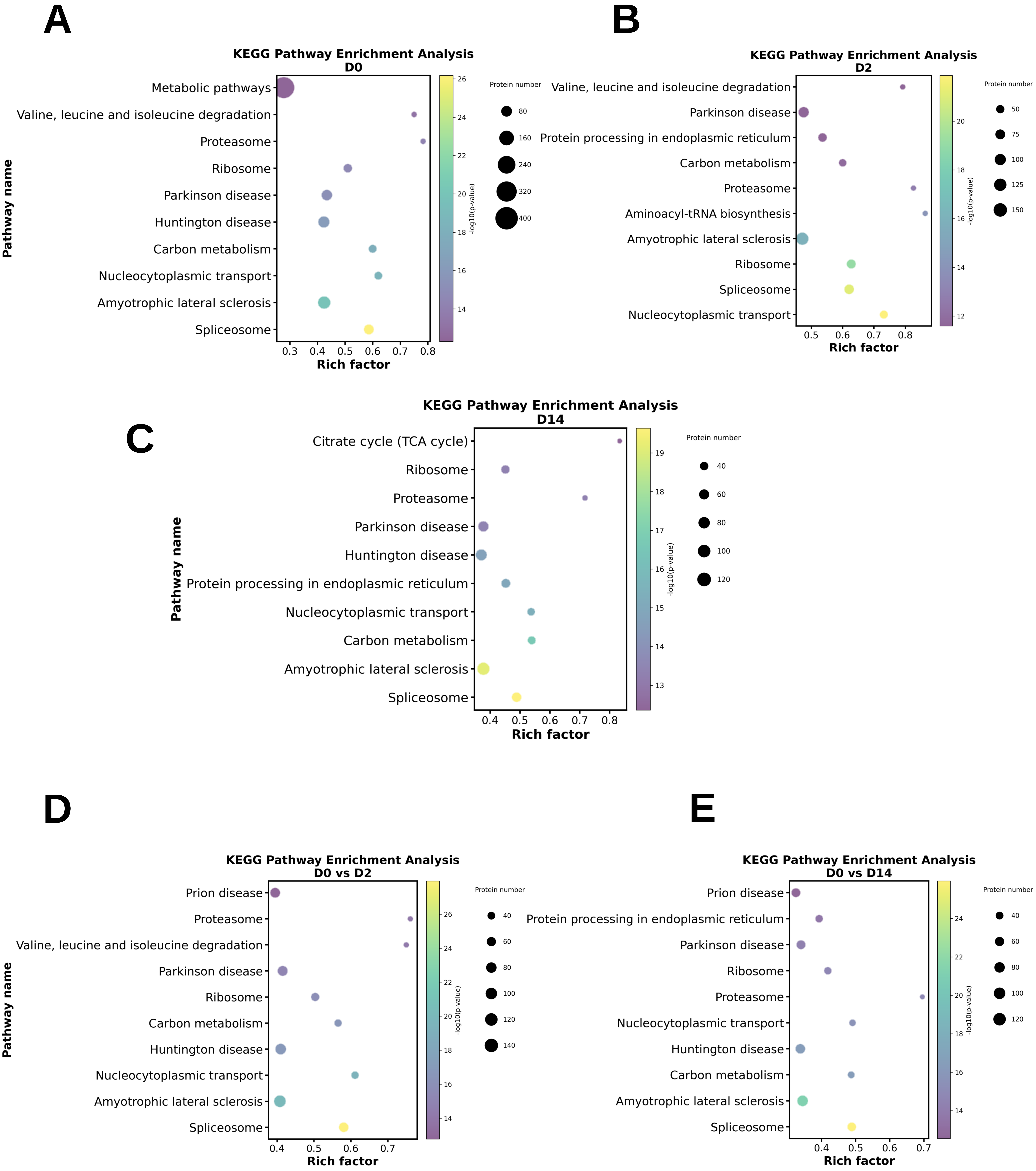


**Figure S19: KEGG enrichment analysis of proteomes from individual experimental groups (A-C), and for D0 vs. D2 or D14 comparisons (D-E) in DIA data.**

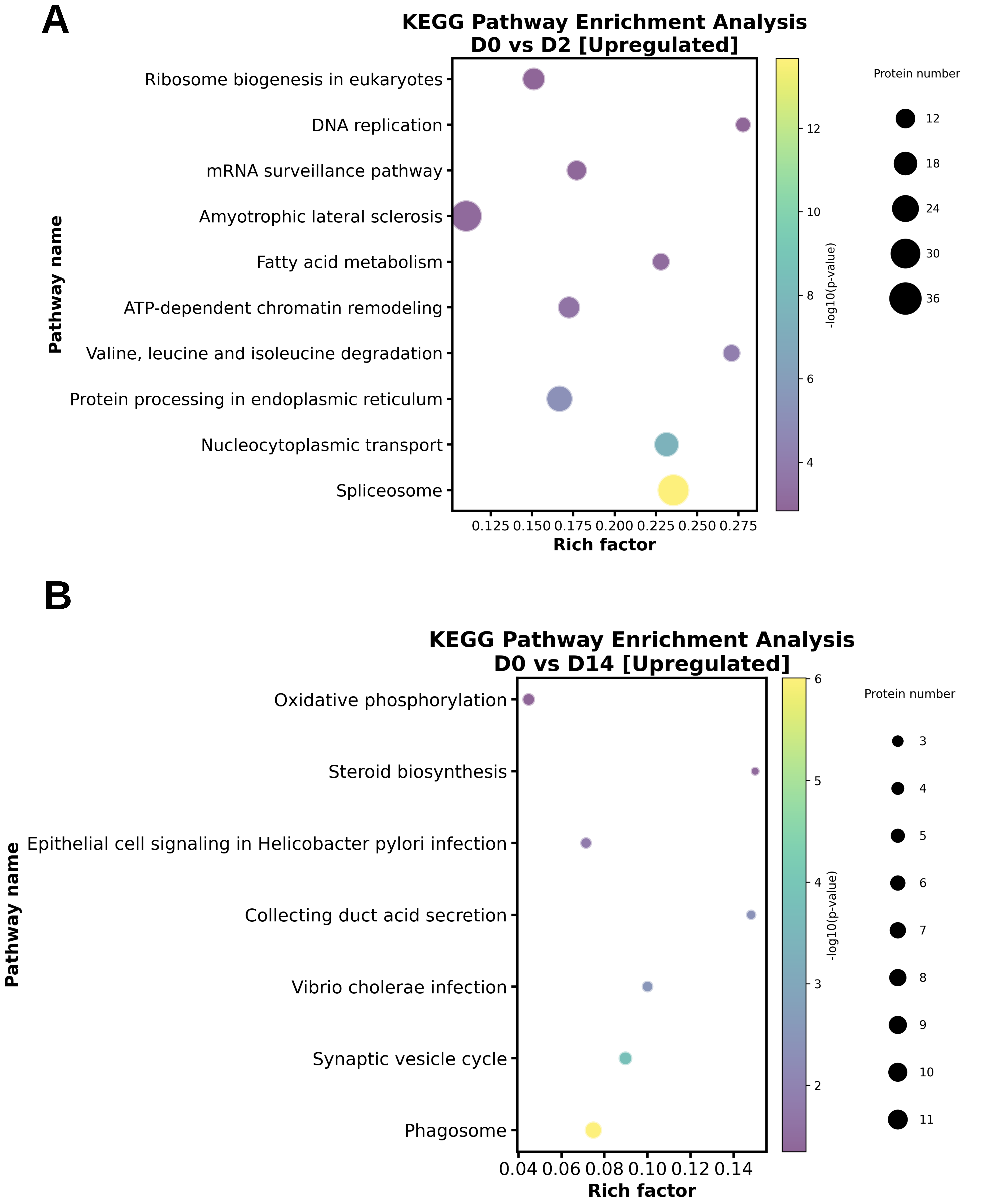


**Figure S20: KEGG enrichment analysis of upregulated proteins in proteomes for A) D0 vs. D2 or B) D0 vs. D14 comparisons from the DIA data.**

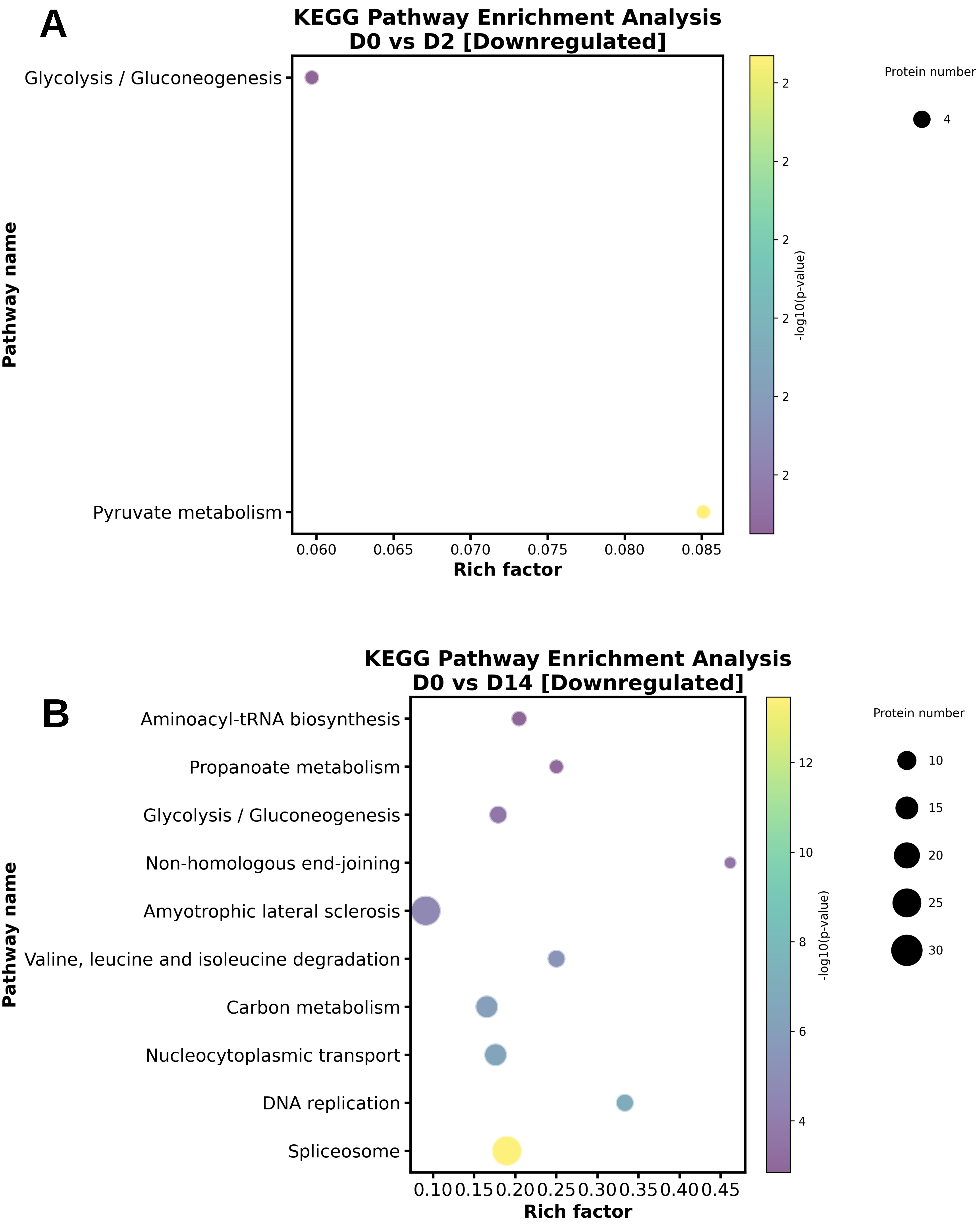


**Figure S21: KEGG enrichment analysis of upregulated proteins in proteomes in A) D0 vs. D2 or B) D0 vs. D14 comparisons from the DIA data.**

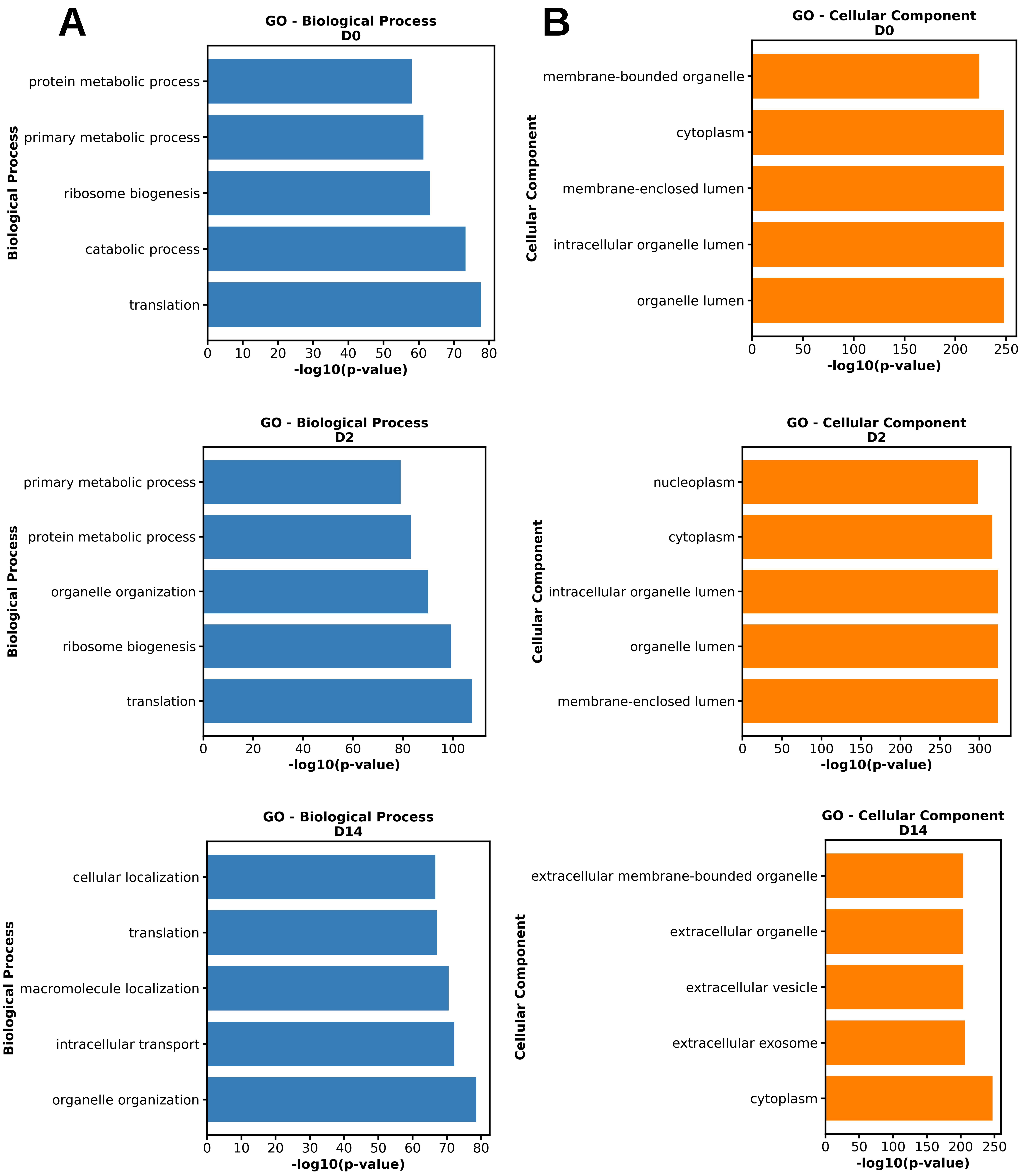


**Figure S22: GO enrichment analysis of proteomes from individual experimental groups in DIA data.** GO enrichment is classified by A) Biological Process, and B) Cellular Component.

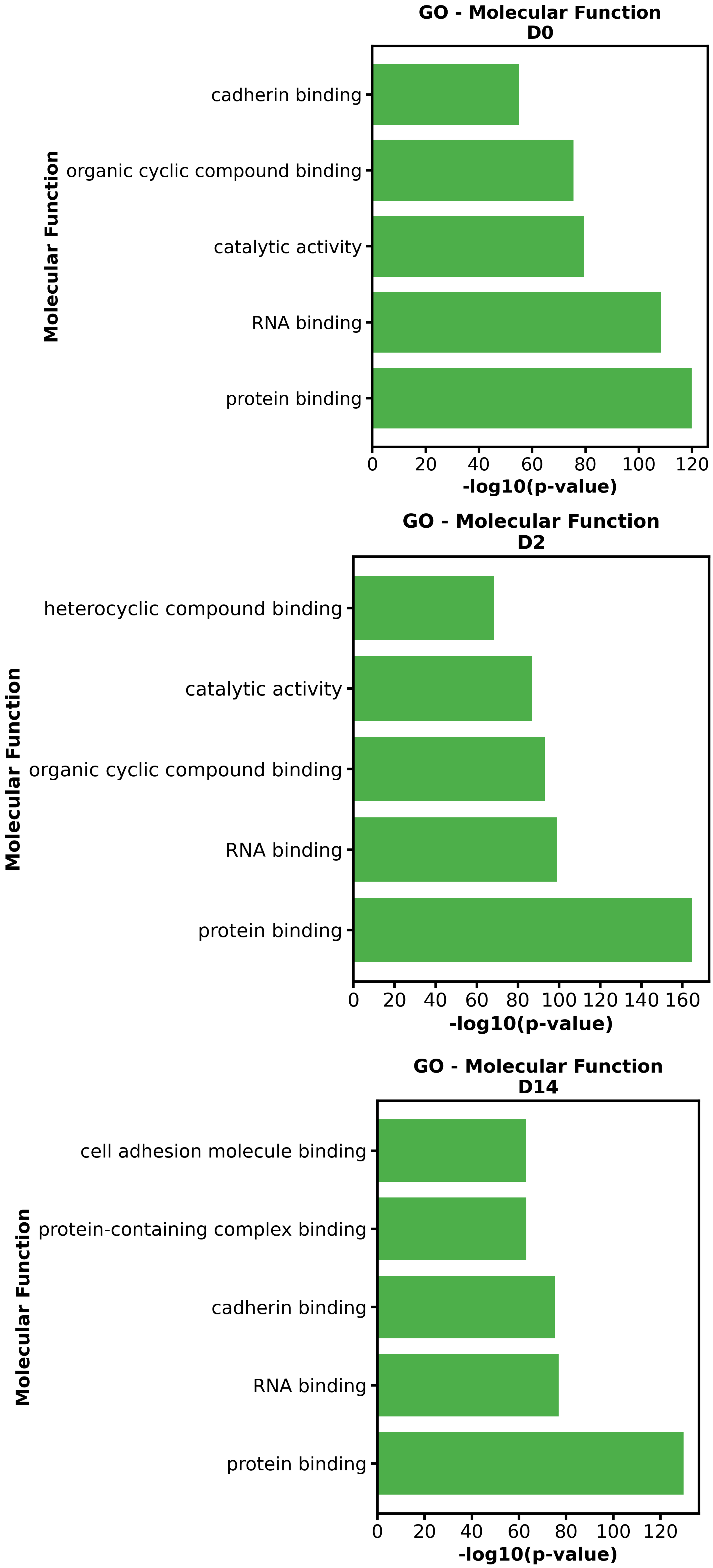


**Figure S23: Molecular function GO enrichment analysis of proteomes from individual experimental groups in the DIA data.**

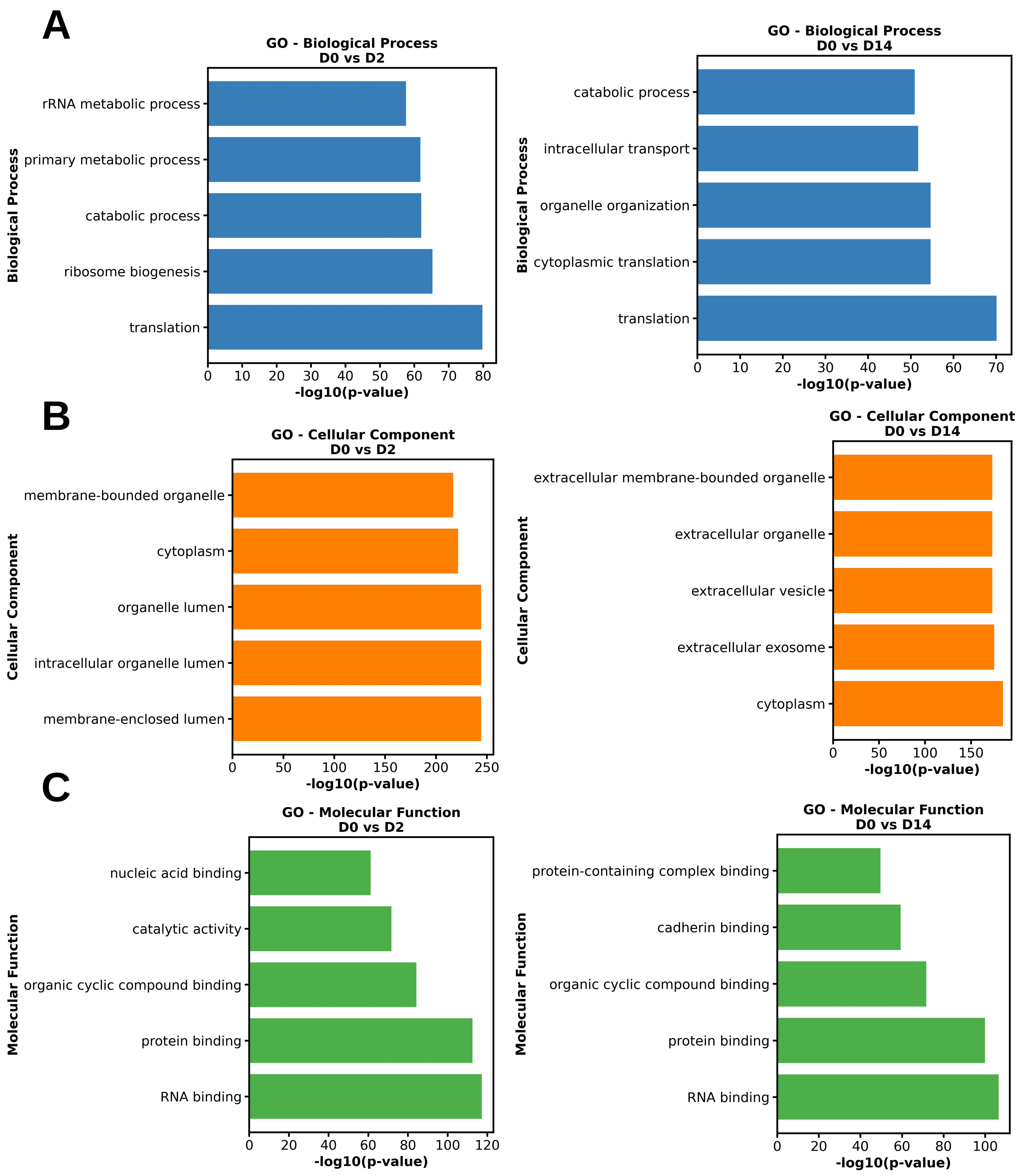


**Figure S24: GO enrichment of proteomes from D0 vs. D2 or D14 comparisons in DIA data.** GO enrichment is classified by A) Biological Process, B) Cellular Component, and C) Molecular Function as depicted in the plots.

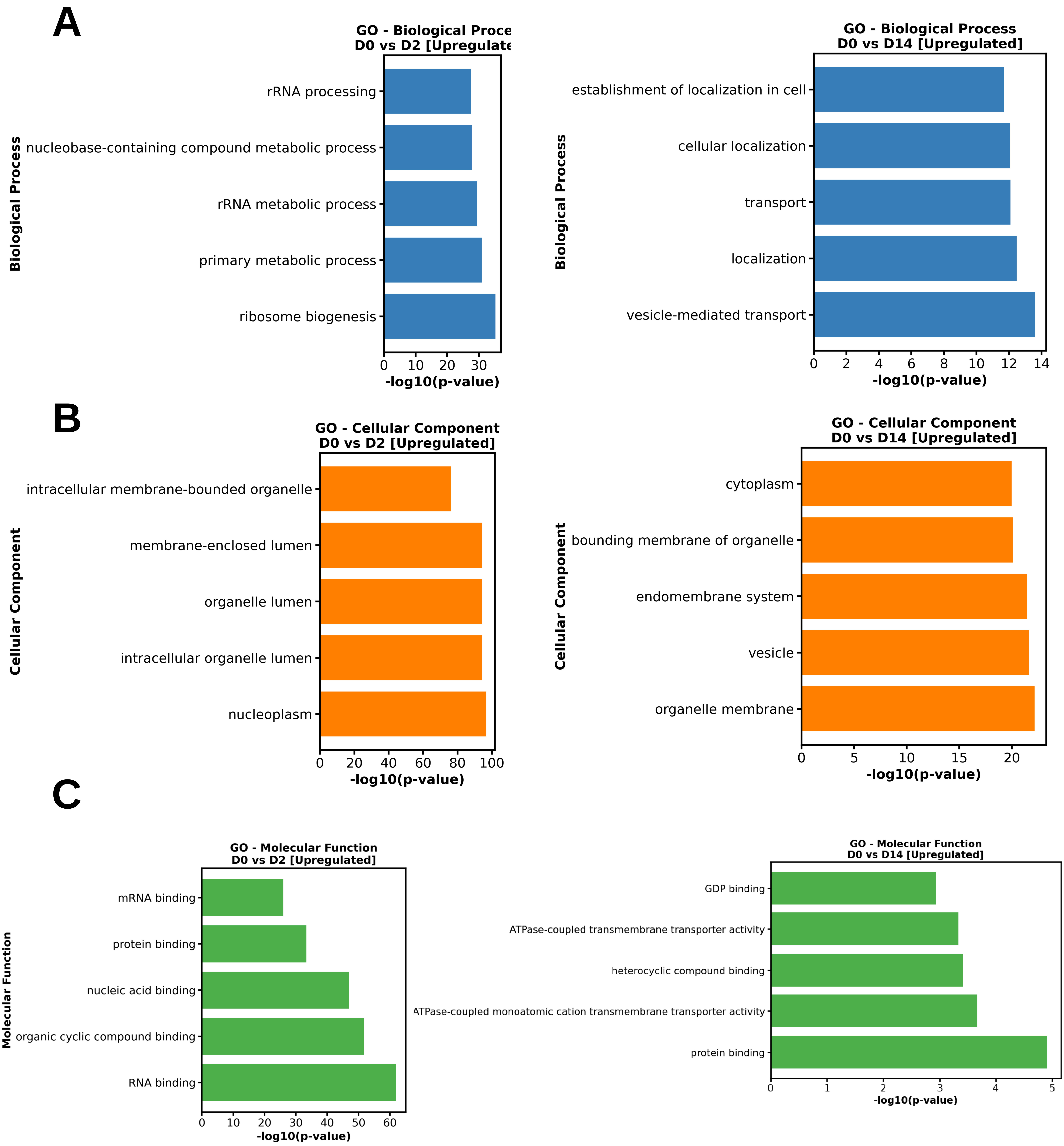


**Figure S25: GO enrichment analysis of upregulated proteins from proteomes in D0 vs. D2 or D14 comparisons from the DIA data.** GO enrichment is classified by A) Biological Process, B) Cellular Component, and C) Molecular Function as depicted in the plots.

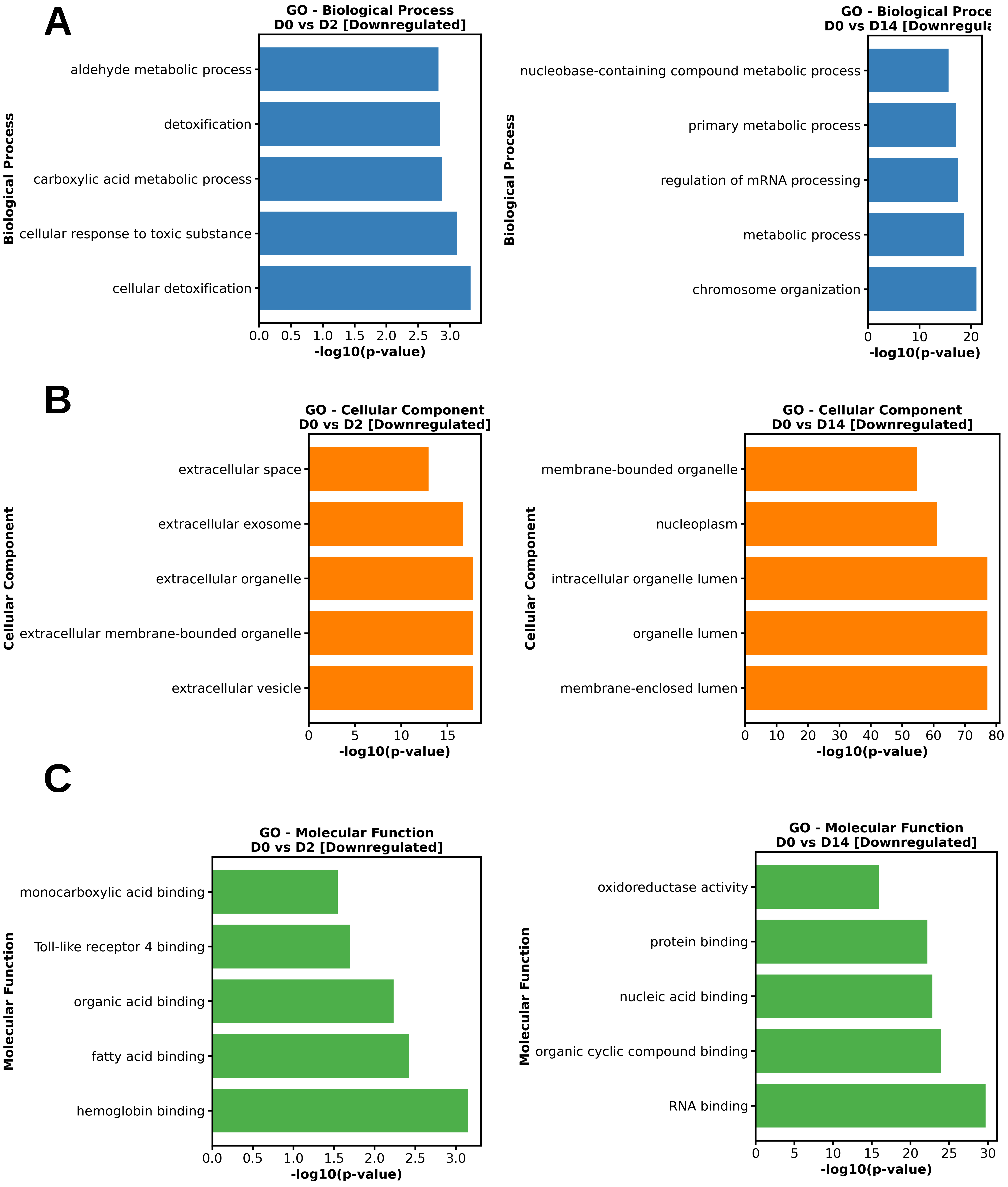


**Figure S26: GO enrichment analysis of downregulated proteins from proteomes in D0 vs. D2 or D14 comparisons from the DIA data.** GO enrichment is classified by A) Biological Process, B) Cellular Component, and C) Molecular Function as depicted in the plots.

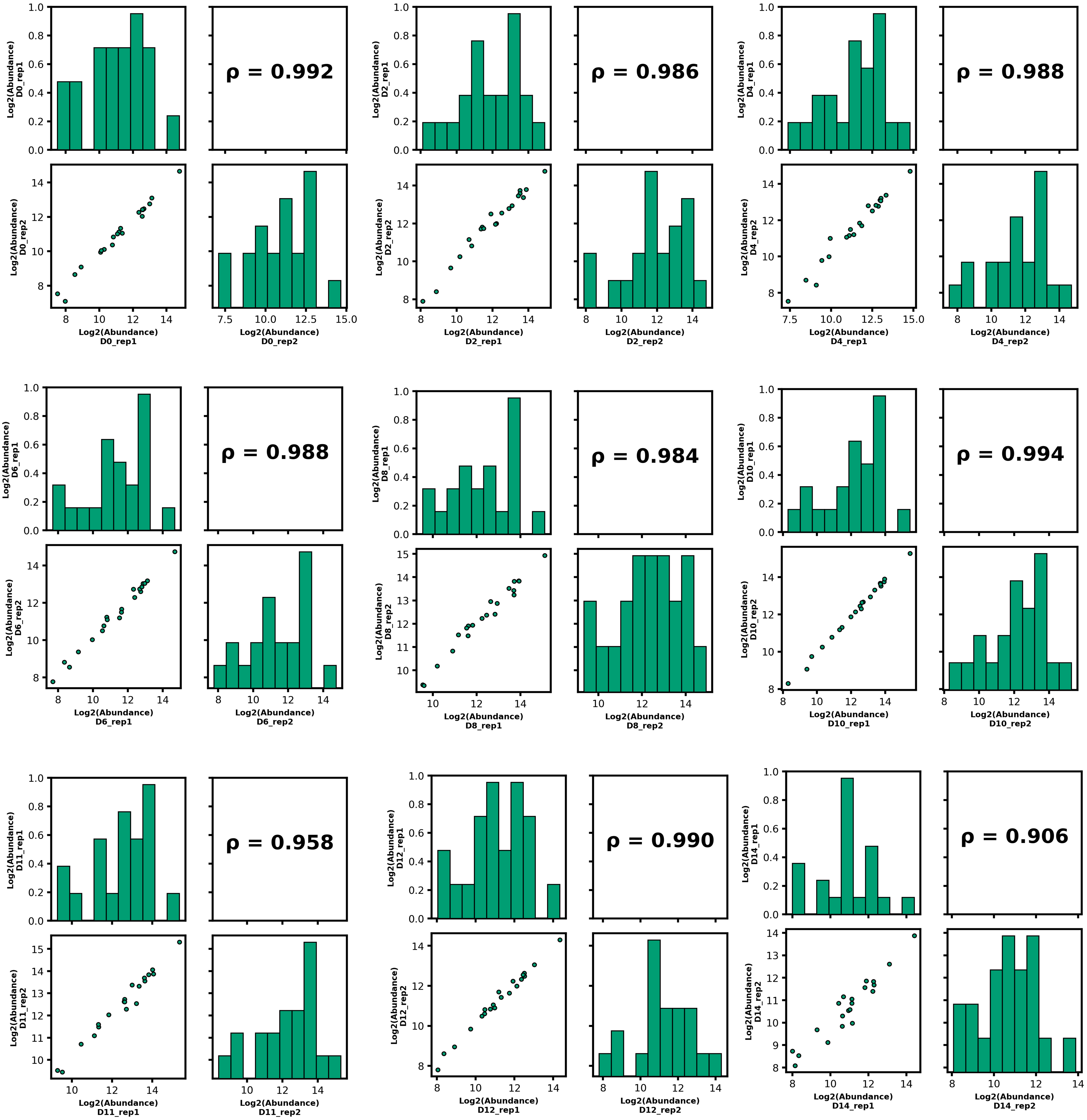


**Figure S27: Spearman’s correlation coefficient (ρ) and data distribution plots among replicates for days, 2, 4, 6, 8, 10, 11, 12, and 14 in PRM data.**

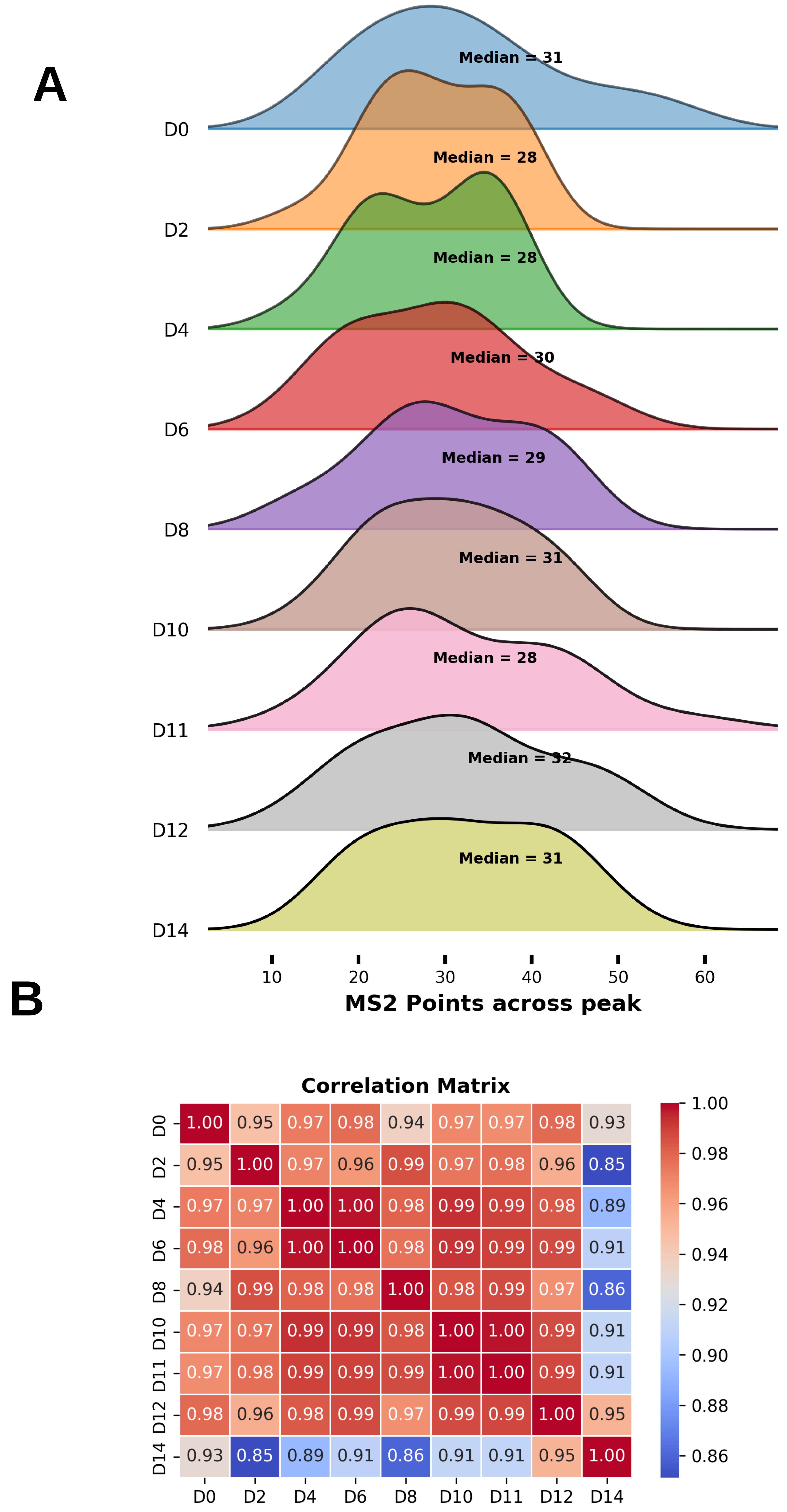


**Figure S28: A) Distribution of MS2 data points across the chromatographic peak and B) correlation matrix among experimental groups from the PRM data.**

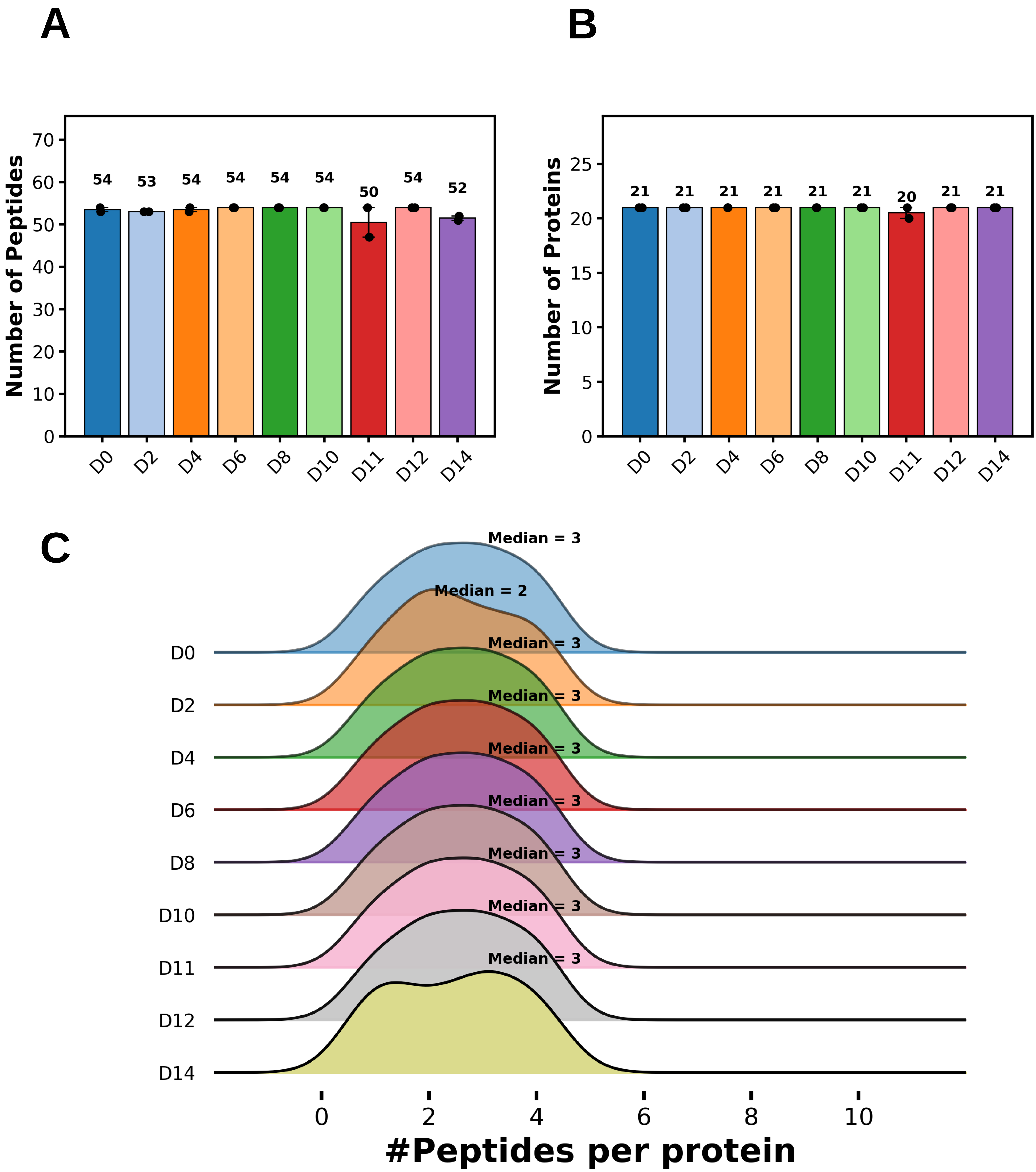


**Figure S29: Number of A) peptides and B) proteins, and C) distribution of number of peptides per protein in PRM data.** Data represent the median ± SD of two biological replicates.

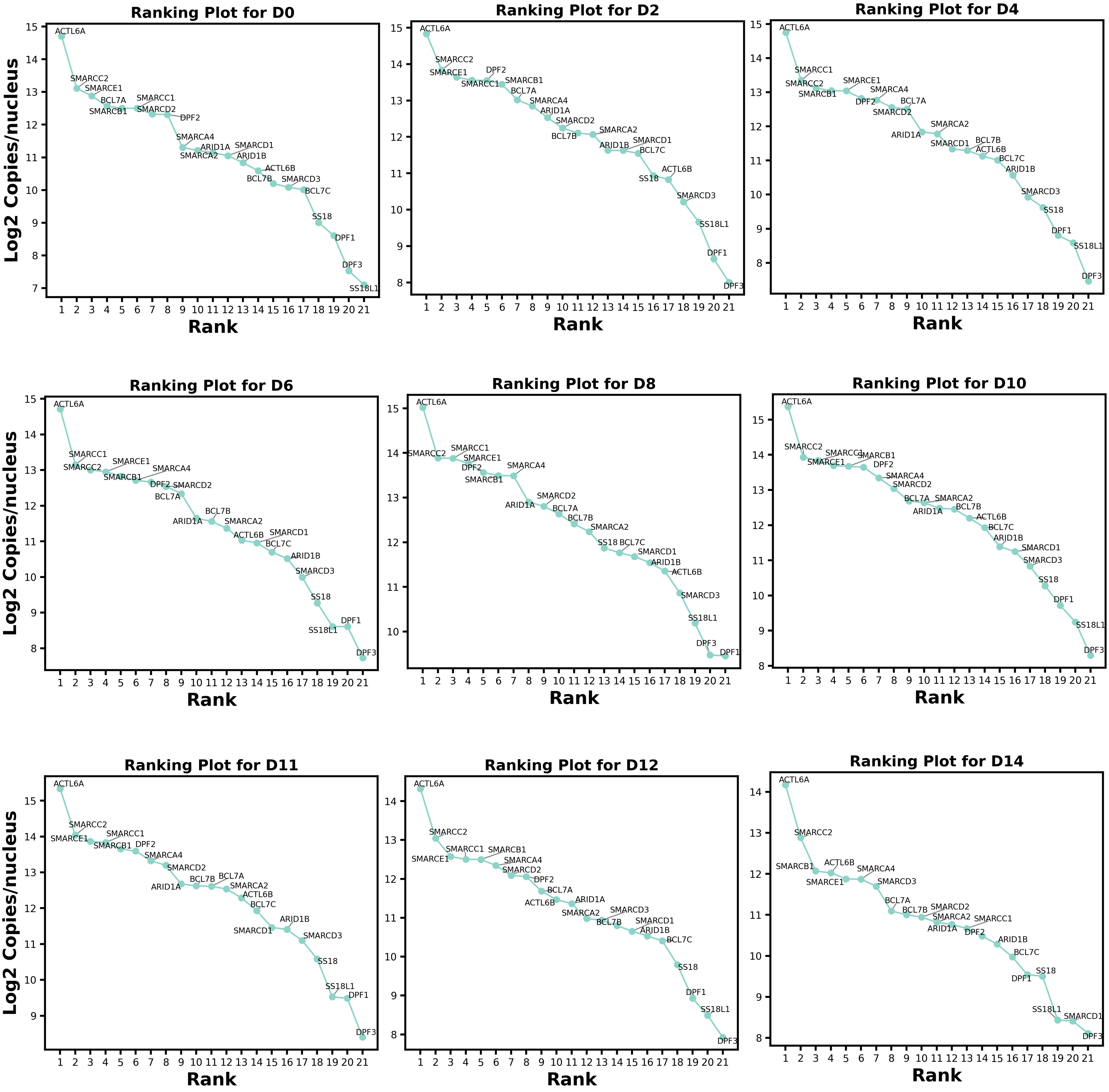


**Figure S30: Protein ranking for erythroid samples collected on days 0, 2, 4, 6, 8, 10, 11, 12, and 14 in PRM data.** The name of every protein in the ranking is depicted in each plot. The Y-axis represents the log2 number of protein copies per nucleus in the proteomes of each experimental group, whereas the X-axis depicts the abundance ranking for proteins in a given proteome.

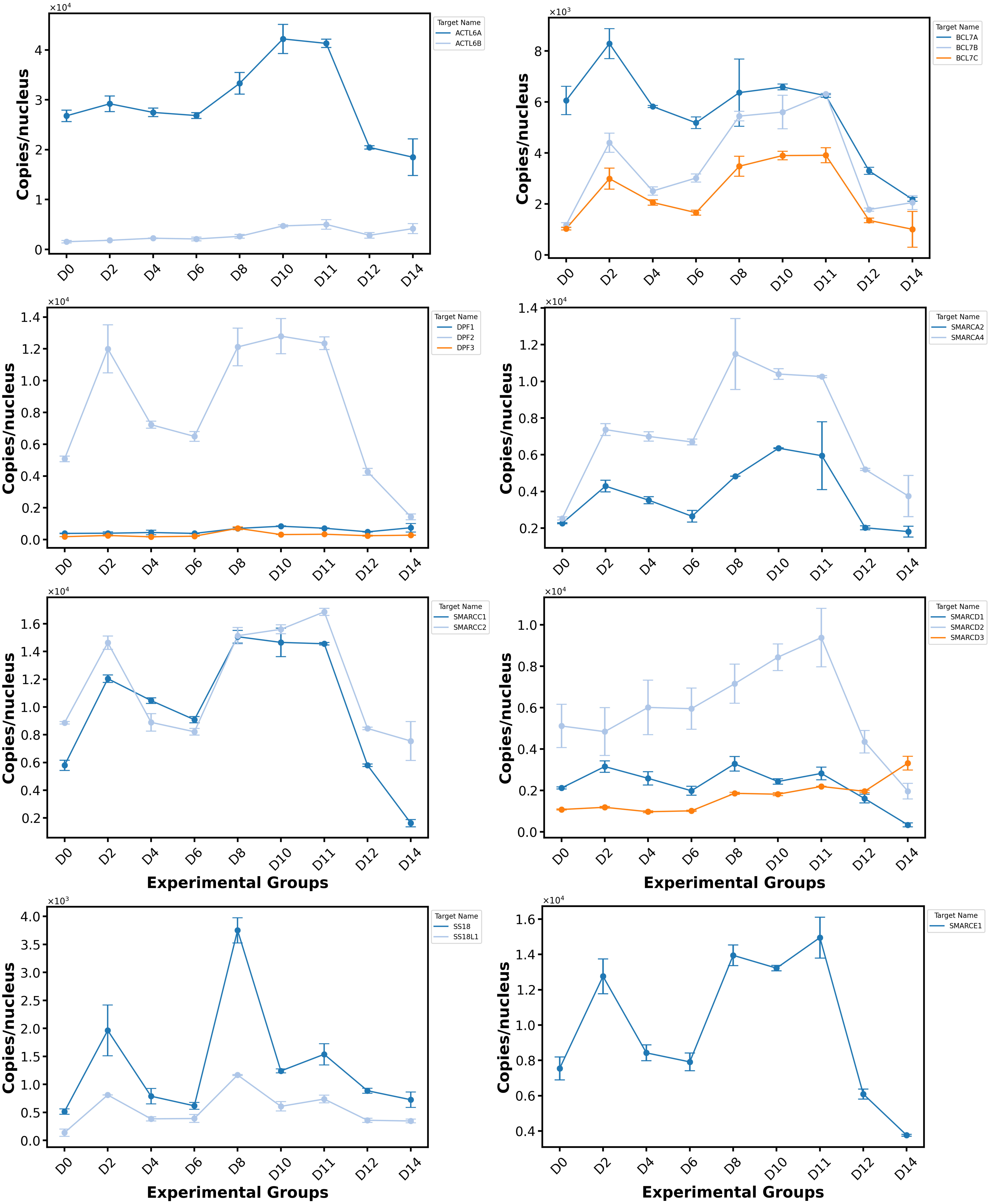


**Figure S31: Trend line plot for protein members of cBAF complex based on the PRM data.** Trend line plot for ACTL6 (ACTL6A, and ACTL6B), BCL7 (BCL7A, BCL7B, and BCL7C), DPF (DPF1-3), SMARCA (SMARCA2, and SMARCA4), SMARCC (SMARCC1, and SMARCC2), SS18 (SS18 AND SS18L1), and SMARCE1 proteins. Data represent the median ± SD of two biological replicates.
